# CD95/Fas ligand mRNA is toxic to cells through more than one mechanism

**DOI:** 10.1101/2022.10.03.510729

**Authors:** Ashley Haluck-Kangas, Madelaine Fink, Elizabeth T. Bartom, Marcus E. Peter

## Abstract

CD95/Fas ligand induces apoptosis through binding of the protein to the CD95 receptor. However, CD95L mRNA also induces toxicity in the absence of CD95. Dying cells exhibit features of DISE (Death Induced by Survival Gene Elimination), a form of cell death mediated by RNA interference (RNAi). DISE relies on targeting mediated by six nucleotides of complementarity between positions 2-7, the 6mer seed sequence of a RISC-bound (R-sRNA), and the 3’UTR of an mRNA, a feature that allows to predict the effect of 6mer seed sequences on cell viability. We now report that CD95L mRNA processing generates an sRNA nearly identical to shL3, a commercial CD95L-targeting shRNA that led to the discovery of DISE. Neither of the miRNA biogenesis proteins Drosha or Dicer are required for CD95L mRNA processing. Interestingly, CD95L toxicity depends on the core component of the RISC, Ago 2, in some cell lines, but not in others. In the HCT116 colon cancer cell line, Ago 1-4 appear to function redundantly in RNAi. In fact, Ago 1/2/3 knockout cells retained sensitivity to CD95L mRNA toxicity. Toxicity was only blocked by mutation of all in-frame start codons in the CD95L ORF. Expression of a toxic CD95L mRNA caused an enrichment for R-sRNAs with toxic 6mer seed sequences, while expression of the nontoxic CD95L mutant enriched for loading of R-sRNAs with nontoxic 6mer seeds. However, CD95L was not the only source of these R-sRNAs. We found that CD95L mRNA may induce DISE directly and indirectly, and that alternate mechanisms may underlie CD95L mRNA processing and toxicity.

## Introduction

CD95/Fas ligand (CD95L) is a well-established inducer of extrinsic apoptosis [1–3]. It is expressed by activated T cells and maintains T cell homeostasis by negatively regulating clonal expansion. In cancer, CD95L expression is induced upon exposure to genotoxic agents, and has been implicated in death by chemotherapeutics and radiation therapy [4, 5]. When membrane bound CD95L binds to its cognate receptor, CD95, it promotes trimerization of CD95 inducing the recruitment of proteins that, in sensitive cells, activate caspases that execute apoptosis [6, 7]. However, we recently reported that overexpression of CD95L induces cell death even in the absence of CD95 receptor [8]. Disrupting engagement of CD95 signaling both by CRISPR/Cas9 knock-out of CD95 and by mutation of CD95L did not block this form of cell death. Treatment with a pan-caspase inhibitor demonstrated that cell death was occurring in a caspase independent fashion exhibiting features of Death Induced by Survival gene Elimination (DISE) [9].

DISE results from the RNA interference (RNAi)-mediated downregulation of networks of genes required for cell survival. Targeting of these essential survival genes depends upon the 6mer seed sequence, nucleotides 2-7, of a short RNA (sRNA) in the RNA-Induced Signaling Complex (RISC). DISE occurs with ROS generation and DNA damage, and exhibits features of apoptosis, autophagy, necroptosis, and mitotic catastrophe [10]. DISE was discovered through experiments using commercial si-/shRNAs designed to target CD95 and CD95L. About 80%of these sequences induced death even in when the target site was deleted [9]. Through a series of experiments we determined that these si-/shRNAs were engaging in a miRNA-like seed-based targeting of hundreds of survival genes. Based on the observation that only six nucleotides (positions 2-7) of the seed sequence were required to mediate this targeting [9], we performed arrayed screens of all 4,096 possible permutations of six nucleotide seed sequences expressed in a non-targeting siRNA backbone in three human and three murine cell lines [11, 12] (6merdb.org). The effect on viability was averaged between cell lines by species and will be referred to as the seed viability.

Based on the data from these seed viability screens, predictions can now be made regarding the effect an sRNA will have on cell viability through 6mer seed-based targeting mediated by the RISC. We developed a gene agnostic small RNA-seq analysis pipeline, called SPOROS, to make these predictions [13]. The pipeline analyzes small RNA-Seq data by tallying reads with the same 6mer seed sequence and attaching the associated seed viability data. Both the overall 6mer seed viability of all sequences or differentially expressed sequences can then be determined. We found that small shifts in the 6mer seed viability of Ago bound sRNAs can predict cellular responses to toxic stimuli [14–16].

We previously reported that CD95L mRNA induces toxicity at least in part through RNAi [8]. In addition to inducing morphological features of DISE, expression of CD95L promoted similar gene expression changes observed in DISE. Sequencing of RISC bound sRNAs (R-sRNAs) revealed the CD95L mRNA was processed to sRNAs that were loaded into the RISC. A similar processing and RISC loading of endogenous mRNAs was also observed with many endogenous mRNAs related to protein translation enriched in the RISC, as determined by a gene ontology analysis. All these R-sRNAs were more abundant in cells lacking the miRNA biogenesis enzyme Drosha, an observation we interpreted as resulting from a global downregulation of miRNAs and thus reduced competition for RISC loading. Interestingly, Drosha k.o. cells and cells lacking another critical miRNA processing enzyme Dicer, were more sensitive to CD95L toxicity, supporting our hypothesis that miRNAs may protect the RISC from the loading of toxic sequences. Knockdown of Ago2, the primary mediator of RNAi, rescued the toxicity. We found that expression of abundant CD95L-derived sequences as siRNAs were toxic to cells. In fact, CD95L-derived sequences also were enriched for toxicity when expressed as shRNAs [17], and this toxicity correlated well with the 6mer seed viability data [11].

In this study we utilize Ago-RNA-pulldown small RNA-Seq combined with a SPOROS analysis (Ago-RP-Seq-SPOROS) to determine the contribution of CD95L mRNA derived sRNAs to the overall 6mer seed viability of the RISC in HCT116 cells lacking various components of the RNAi pathway. We found that the CD95L mRNA is processed into sRNAs that skew more toxic than reads derived from other highly expressed and processed mRNAs. Dicer is not required for the processing of CD95L mRNA nor for the processing of the mRNAs involved in translation. Interestingly, the role of Ago2 in mediating the toxic effects of CD95L mRNA expression appears to be cell type specific. While knockdown of Ago2 in the ovarian cancer cell line HeyA8 blocked toxicity by CD95L mRNA [18], knockdown or knock-out of Ago2 did not rescue DISE induced by either CD95L mRNA or toxic seed containing shRNAs in the colon cancer cell line HCT116. Our data suggests that shifts in the 6mer seed viability of R-sRNAs (derived from CD95L and other genes) affect cell fate. Cell death is associated with an increased loading of sRNAs with toxic 6mer seeds, and decreased loading of sRNAs with non-toxic seeds. This is supported by the expression of various CD95L mutants. Multiple toxic mutants induced a shift towards loading of toxic sRNAs, but a non-toxic mutant exhibited the opposite, increased loading of sRNAs with non-toxic 6mer seeds. Thus, our data suggests that CD95L mRNA can promote DISE directly, through loading of CD95L-derived sRNAs with toxic 6mer seeds, and indirectly, by promoting the loading of other toxic seed containing sRNAs into the RISC.

## Results

### Detection of RISC bound CD95L-derived reads that contain the same sequence as shL3, a commercially available shRNA that is toxic to cancer cells

In our previous studies, one commercial CD95L-derived shRNA was found to be especially toxic and was used in the experiments that led to the discovery of DISE [17]. We referred to this sequence as shL3. In an experiment to determine the mechanism of shL3 toxicity, we expressed shL3 in 293T cells in which the shL3 target site in CD95L was deleted [17]. Sequencing of the total sRNA revealed that the 5-p arm, or sense strand, of the shRNA was predominantly expressed. In fact, there were 10 times more reads derived from the shL3 sense strand (5,039,726 sense reads to 477,308 antisense reads, **Fig. S1A**). Of the 5 million shL3 sense reads, 85.6% shared the same 6mer seed sequence, **ACTGGG** (red box in **Fig. S1B**). Based on our 6mer seed viability screens, we predict the average viability associated with this R-sRNA to be 37.6%. Alternative processing of this sequence (seen in 2.8% of cases) resulted in the generation of a seed that is even more toxic (GACTGG, seed viability 28%) (dark grey arrowhead on 5p arm, **Fig. S1B**). Conversely, the less abundant antisense strand was predicted to give rise to nontoxic sRNAs and the main species (full arrow head on 3-p arm, **Fig. S1B**) is expected to have a 6mer seed viability of 80% (seed: ACAAAG, green box in **Fig. S1B**). Thus, we concluded that the toxicity we observed from shL3 expression resulted from the activity of the much more abundant sense strand of the shL3-derived sequence.

Previously, we confirmed that CD95L-derived R-sRNAs exerted toxicity when expressed as siRNAs (Figure 4S1B in [8]). Among the most toxic of these abundant sequences was cluster #15 (cl15). Closer inspection revealed that cl15 is nearly identical in sequence to the shL3 sense strand. We have now reanalyzed the Ago-RP-Seq data from these HCT116 Drosha k.o. cells expressing pLenti-CD95L NP, a mutant cDNA that does not produce full length CD95L protein and has a point mutation that prevents the truncated protein from binding to CD95 [8]. We found that ~10% of reads derived from CD95L contain the toxic shL3 sequence (**Fig. 1A, B**). In fact, one of the two predominant species in the RISC, cl15.2, is nearly identical to shL3 (only position 1 differs, as in shL3 it was derived from the pTIP vector sequence). Plotting the number of CD95L reads against their associated 6mer seed viabilities revealed that cluster 15 is a substantial source of toxic sRNAs derived from CD95L (**Fig. 1C**). We confirmed that the sequence cl15.2 exerts a negative growth effect through RNAi as HCT116 Drosha k.o. cells were more sensitive than wild-type (wt) cells to the toxicity when the sequence was introduced as an siRNA (**Fig. 1D**, left and center panel). This was in agreement with previous observations that Drosha k.o. cells are more sensitive to other toxic siRNAs, such as the CD95L derived siRNA, siL3 [10], and an siRNA carrying the toxic consensus seed, GGGGGC [12]. In contrast, Ago 1/2/3 k.o. cells were completely resistant to cl15.2 siRNA toxicity (**Fig. 1D**, right panel). These data suggest that CD95L can give rise to an R-sRNA that is almost identical to the commercial toxic CD95L derived shRNA that resulted in the discovery of DISE.

**Figure 1.**
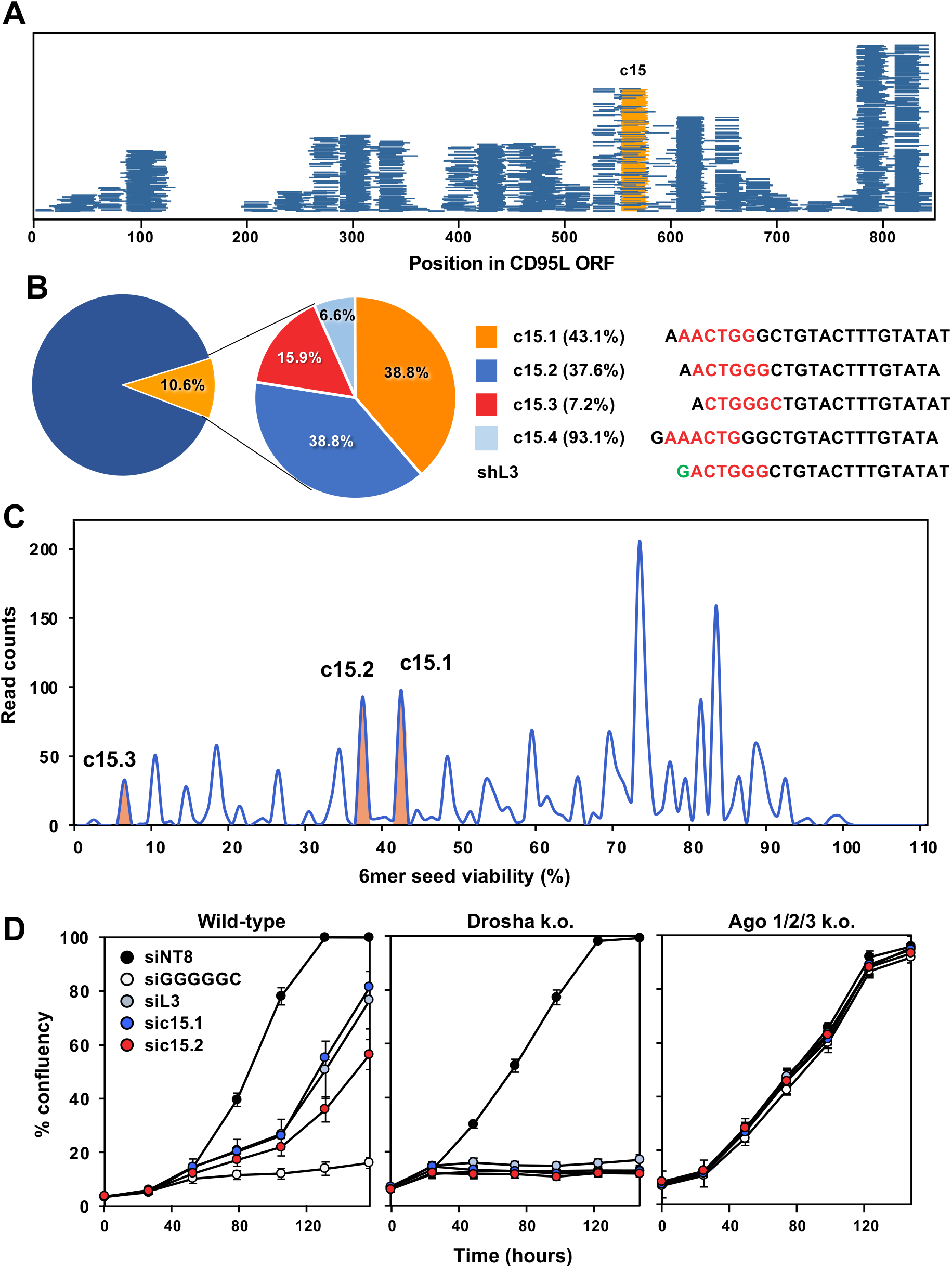
CD95L mRNA is processed to generate a sequence analogous to the toxic shRNA, shL3. **(A)** R-sRNAs derived from CD95L in HCT116 Drosha k.o. cells infected with pLenti-CD95L NP mapped along the ORF. Reads with the same sequence and 6mer seed as shL3 are indicated in orange. **(B)***Left*, % of R-sRNAs derived from CD95L cluster 15. *Right*, breakdown of the percent of reads in cluster 15 (left) by 6mer seed (red lettering) and predicted 6mer seed viability (in parentheses). **(C)** Seed viability graph of CD95L-derived sequences of Drosha k.o. cells infected as in A with seed viability represented on the x-axis and normalized read counts on the y-axis. The three main read peaks that correspond to c15 are labeled in orange. **(D)** Confluency over time of wt, Drosha k.o., or Ago 1/2/3 k.o. cells transfected with 10 nM of control siRNA siNT8, the toxic human consensus containing siRNA siGGGGGC, the CD95L derived DISE inducing siRNA siL3, or siRNAs corresponding to c15.1 and c15.2. Data are representative of two independent experiments.

**Figure 2.**
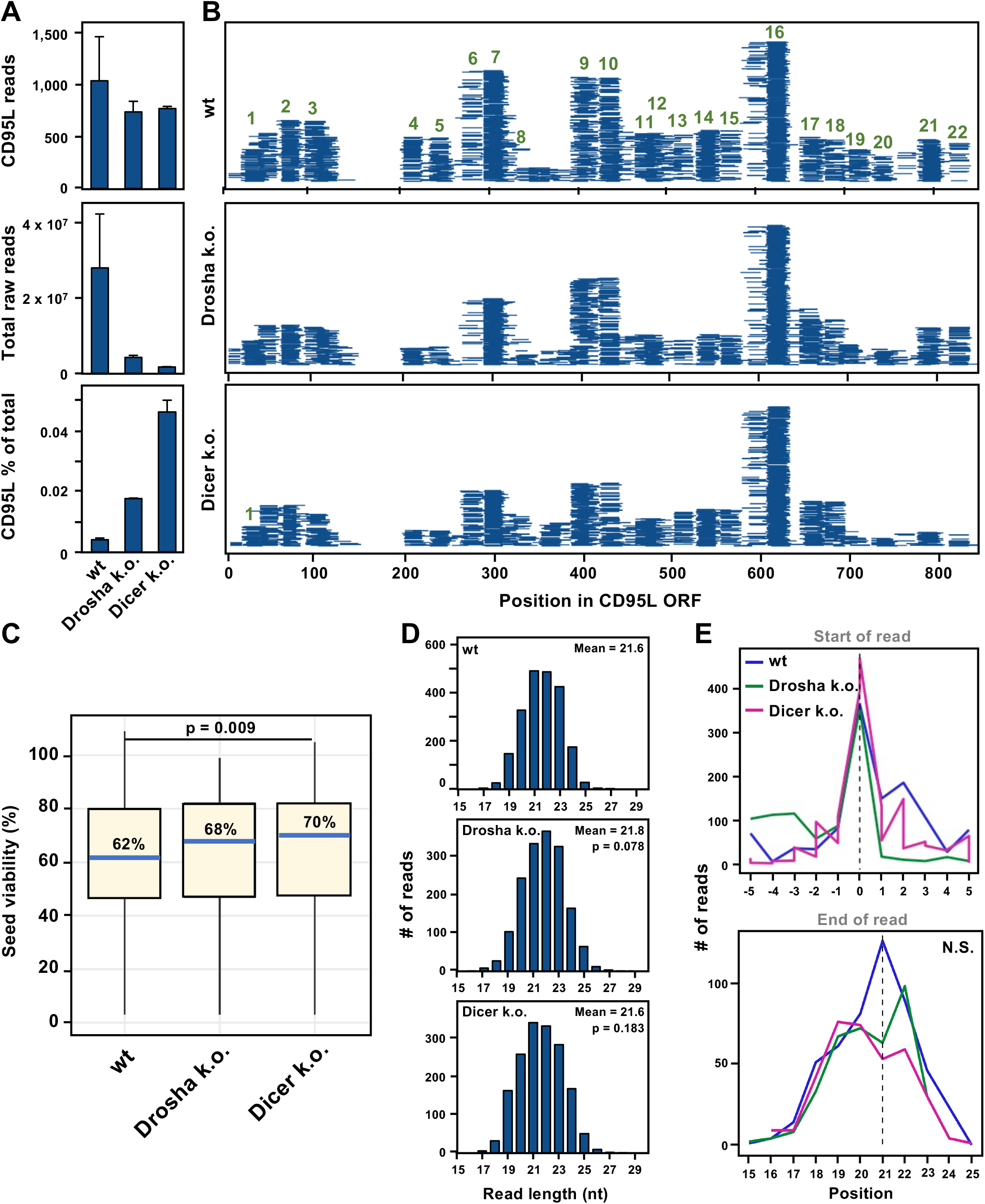
Loss of Dicer has no effect on the processing of CD95L-derived R-sRNAs. **(A)** Wild-type, Drosha k.o., or Dicer k.o. HCT116 cells were infected with pLenti-CD95L NP and Ago-RP-Seq analysis was performed 100 hrs after infection. *Top*, Raw R-sRNA counts aligning to CD95 NP. *Center*, the total raw R-sRNA reads sequenced per sample. *Bottom*, percentage of total raw R-sRNA reads derived from CD95L NP. Averages of two replicates are shown. Error bars represent the variance of the mean. **(B)** Mapping of CD95L NP-derived sRNAs along the CD95L ORF. Each horizontal line represents one read. A combination of reads from both replicates are displayed. **(C)** Box plots representing the distribution of the 6mer seed viability associated with CD95L-derived reads in the RISC in the samples in A. Black lines and labels represent the median 6mer Seed viability in each sample. The Kruskal-Wallis test p-value is given. **(D)** Bar plots represent the read length distribution of all CD95L reads in the RISC. Significance was determined using the Kolmogorov-Smirnov test with Bonferroni corrected p-values. **(E)** The 10 most abundant stacks from (B) were analyzed for differential trimming. *Top*, the 5’ start position of the most abundant reads in each stack is indicated at 0. Reads with 5’ start sites either 5 nucleotides upstream or downstream were tallied. *Bottom*, the 3’ stop site of the 10 most abundant reads were tallied. Kruskal-Wallis p-value = 0.771 (N.S., not significant).

**Figure 3.**
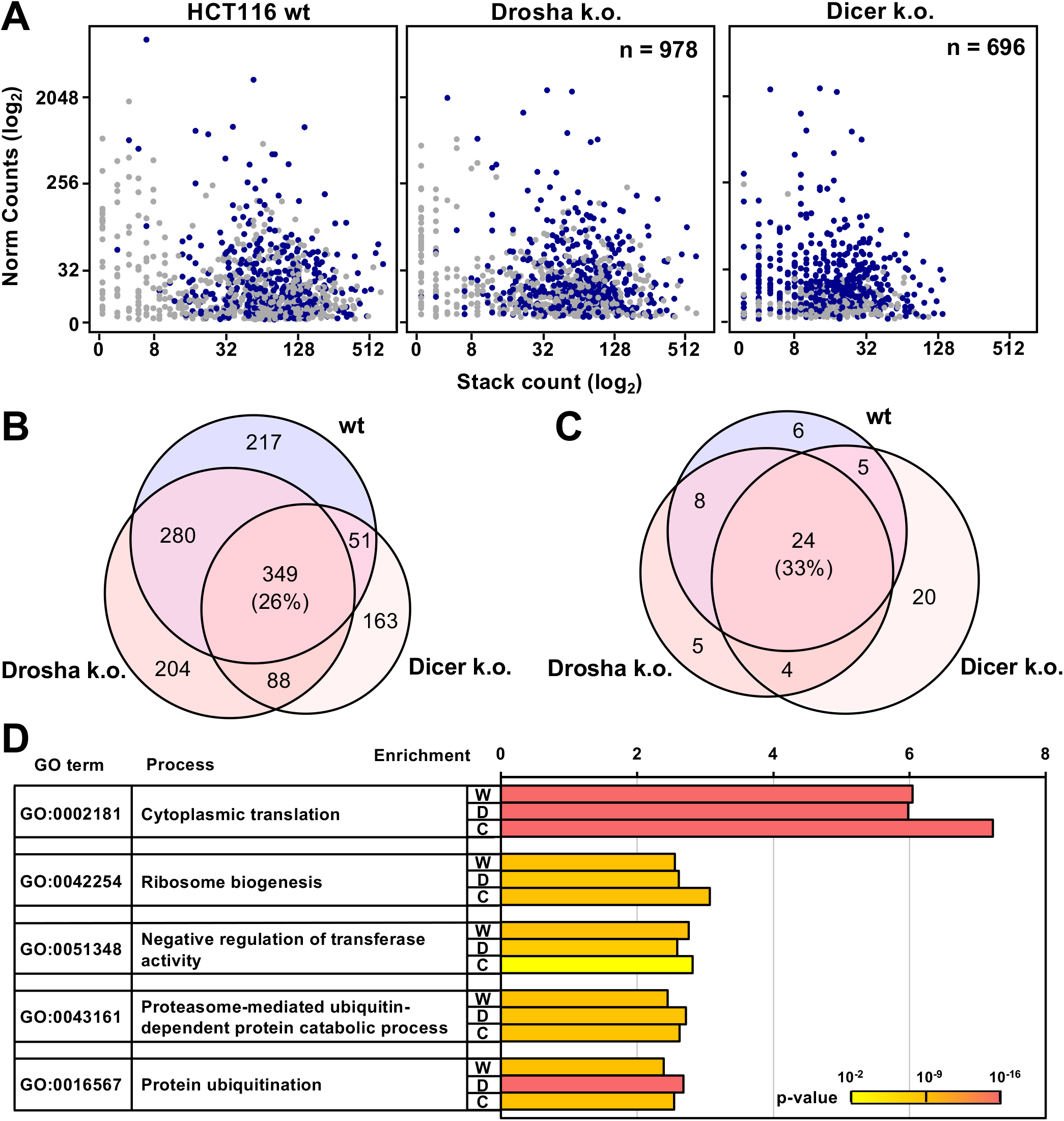
Protein-coding mRNAs involved in cytoplasmic protein translation are processed and loaded into the RISC independent of Dicer. **(A)** Dot plots representing processed mRNAs found in the RISC of HCT116, Drosha k.o., and Dicer k.o. cells. Normalized gene expression is represented on the y-axis and stack count on the x-axis. A stack is defined as at least 10 raw reads mapping to a mRNA transcript with the same 5’ start site. Shared genes between all datasets are in dark blue. **(B)** Venn diagram displaying the overlap of genes processed in, Drosha k.o., and Dicer k.o. cells. **(C)** Overlap of significant GO terms (DAVID, Bonferroni adj. p-value < 0.05) enriched at least 1.5 fold in the three genotypes. **(D)** The top five enriched GO terms in all three genotypes. The Bonferroni corrected p-values are represented by color from p= 0.002 in yellow to p = 1.9 x 10^-16^ in pink.

**Figure 4.**
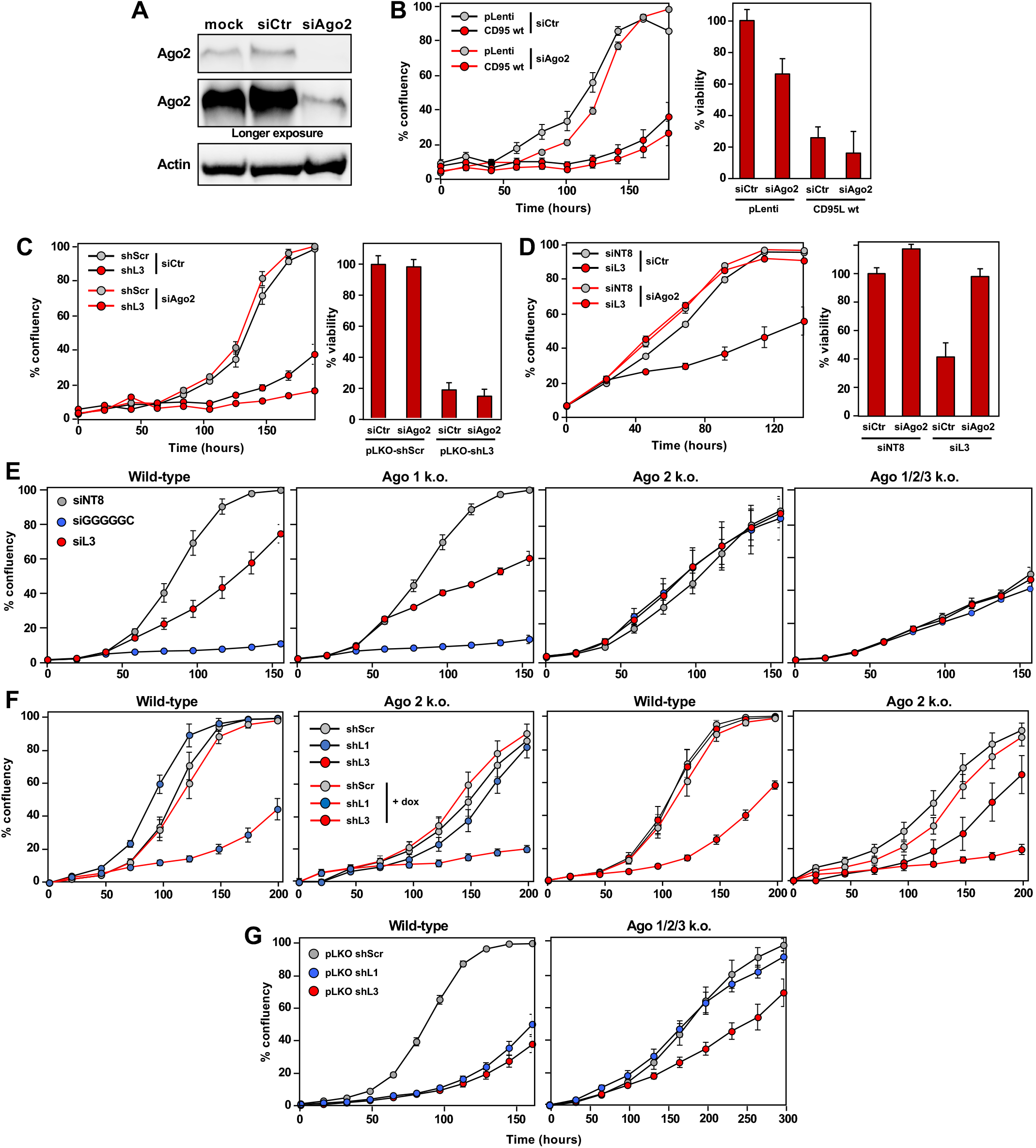
Ago2 is not required for toxicity of CD95L mRNA or DISE-inducing si-/shRNAs in HCT116 cells. **(A**) Western blot analysis of Ago2 expression in HCT116 Drosha CD95 d.k.o. c12 mock transfected or transfected with 25 nM siAgo2 or scrambled control (siCtr) siRNA SmartPools. **(B)***Left*, Percent confluency over time of HCT116 Drosha CD95 d.k.o. c12 transfected with either 25 nM siAgo2 or siCtr siRNAs, and subsequently infected with pLenti or pLenti-CD95L *Right*, relative cell viability at 120 hours. **(C)***Left*, percent confluency over time of HCT116 Drosha CD95 d.k.o. c12 transfected with either 25 nM siAgo2 or siCtr siRNAs, and subsequently infected with pLKO-shScr or pLKO-shL3 (left). *Right*, relative cell viability at 120 hours. **(D)***Left*, Percent confluency over time of HCT116 Drosha CD95 d.k.o. c12 transfected with either 25 nM siAgo2 or siCtr siRNAs, and subsequently transfected 10 nM siL3 or siNT8 non-targeting control. *Right*, relative cell viability at 96 hours. **(E)** Percent confluency over time of HCT116 (left), HCT116 Ago1 k.o. (center left), HCT116 Ago2 k.o. (center right), and HCT116 Ago 1/2/3 k.o. (right) transfected with 10 nM siGGGGGC, siL3, or siNT8 control. **(F)** Percent confluency over time of HCT116 wt and HCT116 Ago2 k.o. cells stably expressing either doxycycline (dox) inducible pTIP-shScr, pTIP-shL1, or pTIP-shL3. Cells were left untreated or treated with 100 ng/ml dox at 0 hours. **(G)** Percent confluency over time of HCT116 wt (left) and HCT116 Ago 1/2/3 k.o. cells (right) infected with pLKO-shL1, pLKO-shL3 or pLKO-shScr. Error bars in B-D represent the standard deviation of quadruplicates.

### Processing of CD95L mRNA is independent of Dicer

In our previous analysis, many of the CD95L-derived R-sRNAs were found to localize to double stranded regions in the secondary structure of CD95L [8]. Dicer has been reported to bind to mRNAs [19], and may even cleave mRNA substrates [20]. We therefore wondered whether Dicer was involved in the processing of CD95L mRNA. In our previous report, we had detected some CD95L-derived sRNAs in Dicer k.o. cells using real-time quantitative (q)PCR, suggesting that Dicer may not be required for the processing of CD95L mRNA [8]. To test whether this finding could be generalized and to determine if either the pattern of CD95L processing or the loading of the derived sRNAs into the RISC was dependent on Dicer, we infected HCT116 wt, Drosha k.o. and Dicer k.o. cells with pLenti CD95L NP and sequenced the R-sRNAs in these cells. As previously reported, we observed greater toxicity in Drosha and Dicer k.o. cells expressing CD95L, both by a greater reduction in cell growth and cell viability (**Fig. S2A, B**). Interestingly, CD95L NP was also more highly expressed at the mRNA level (**Fig. S2C**).

Analysis of R-sRNAs in HCT116 Dicer k.o. cells revealed that, like Drosha k.o. cells, a variety of sRNAs other than miRNAs are found in the RISC. While the RISC of HCT116 wt cells mostly consisted of miRNAs, Drosha and Dicer k.o. cells, unable to produce most mature miRNAs, instead had a variety of other RNA species bound to the RISC (**Fig. S3A**). An analysis of the top five most abundant R-sRNAs in each genotype revealed that miRNAs that dropped in abundance in k.o. cells (**Fig. S3A, B)**, and were replaced by many other sRNA species in Drosha k.o. and Dicer k.o. cells (**Fig. S3A, C, D**). It was surprising that the predominant R-sRNAs in Dicer k.o. cells were derived from tRNAs fragments (**Fig. S3A, D**) as Dicer has been demonstrated to process some tRNAs [21]. This data likely reflects the fact that RNAs similar in structure and function can be processed via different mechanisms.

To determine if Dicer is involved in the processing of CD95L mRNA, we identified all RISC bound raw reads in our samples that uniquely mapped to the CD95L ORF (**Fig. 2A, B**). In this experiment, we found about the same number of reads derived from CD95L in all three genotypes (**Fig. 2A**, top). However, when we accounted for the number of reads in each sample (**Fig. 2A**, middle), we noticed that as we previously observed, more of the RISC was occupied by CD95L-derived sRNA in Drosha k.o. cells (**Fig. 2A**, bottom), and surprisingly, an even higher percentage of the RISC was occupied by CD95L reads in Dicer k.o. cells. Examining the pattern of processing revealed no major differences between genotypes (**Fig. 2B**). Most reads seemed to be derived from similar regions of the mRNA. However, an analysis of the predicted 6mer seed viability of CD95L reads revealed that CD95L reads in Dicer k.o. cells trended significantly less toxic than in wt cells (**Fig. 2C**), which could suggest some differential processing at the 5’ end of the sRNA. However, the lengths of CD95L reads did not vary significantly between genotypes (**Fig. 2D**). A closer examination of the 5’ start site and 3’ end site of the most abundant reads revealed that the processing of CD95L was remarkably similar between genotypes. Thus, it is not likely that Drosha or Dicer function in the processing of CD95L mRNA.

### RISC bound reads of highly processed mRNAs in both Drosha and Dicer k.o. cells are derived from genes that function in protein translation

In our previous report, we found that CD95L was not the only protein coding gene processed and loaded into the RISC. Many mRNAs involved in translation and the cell cycle were processed in a similar manner, particularly in Drosha k.o. cells [18]. To confirm this finding and determine if these mRNAs were processed in both Drosha and Dicer k.o. cells, all R-sRNAs from cells infected with pLenti CD95L NP were aligned to the human genome and reads mapping uniquely to protein coding genes were plotted (**Fig. 3A**). Genes were considered processed if there were three or more stacks mapping to a gene and at least 10 normalized reads. These criteria were met by many hundreds of genes in each genotype (**Fig. 3A**). Of these, 349 (26%) processed mRNAs were shared (**Fig. 3B**). To determine if the processed mRNAs were derived from genes with similar functions, we performed a gene ontology (GO) analysis, and found that 33% of enriched GO terms were shared (**Fig. 3C**). Consistent with our previous report, genes involved in protein translation were highly enriched (**Fig. 3D**). Other GO terms that were shared were consistent with the Drosha and Dicer independently processed genes being involved in proteasome mediated protein degradation.

Examining the pattern of processing in mRNA transcripts with many stacks revealed subtle differences in the nature of the reads loaded into the RISC of HCT116 wt, Drosha, and Dicer k.o. cells (**Fig. S4**). While processing of FAT1 was very similar between genotypes (**Fig. S4A**), ACTG1 exhibited some differences in processing and/or loading of reads into the RISC (**Fig. S4B**). In other genes certain read stacks abundant in Drosha k.o. and wt cells were almost absent in Dicer k.o. cells (red arrows in **Fig. S4C, D**). Nevertheless, our data do not support a role for Dicer in the processing and loading of these sequences. Mapping abundant R-sRNAs derived from EEF1A1 to the predicted secondary structure revealed that they did not map to stem loop structures expected for potential Dicer substrates (data not shown). This is consistent with our processing analysis of CD95L mRNA. It is not likely that Dicer is a mediator of the processing of these mRNAs. However, there remains a possibility that Dicer could play a role in the loading of the mRNA-derived sRNAs into the RISC.

### In HCT116 cells the toxicity of neither CD95L mRNA nor CD95L-derived shRNAs depends on Ago2

We previously showed that Ago2 was required for CD95L toxicity in HeyA8 CD95 k.o. cells [8]. Thus, we were wondering whether the catalytic activity of Ago2 could be involved in the trimming of CD95L reads in addition to mediating CD95L toxicity through RNAi. To test this hypothesis, we generated HCT116 Drosha CD95 double knock-out (d.k.o.) cells. These cells would allow for analysis of wt CD95L mRNA processing while preventing the apoptosis inducing activity of any CD95L protein expression. Three homozygous CD95 mutants were isolated (**Fig. S5A**) all three did not express any detectable CD95 protein (**Fig. S5B**) and retained sensitivity to pLKO-shL3 (**Fig. S5C, D**). Efficient Ago2 knockdown was achieved in HCT116 Drosha CD95 d.k.o. clone 12 (**Fig. 4A**). However, this did not rescue CD95L toxicity (**Fig. 4B**). This was not a clonal effect as Ago2 k.d. also did not rescue toxicity induced by CD95L NP in HCT116 wt cells or parental Drosha k.o. cells (**Fig. S6A**). It is possible that the exclusive role of Ago2 in CD95L toxicity is cell type specific; Knockdown of Ago2 was able to rescue toxicity in HeyA8 cells [18].

Our data led us to wonder if knockdown of Ago2 was sufficient to block all RNAi in HCT116 cells. Interestingly, knockdown of Ago2 in the same d.k.o. clone was not sufficient to block DISE induced by shL3 (**Fig. 4C**). This was surprising, as Ago2 is thought to be the primary mediator of RNAi. To determine if this effect was specific to shL3, we transfected toxic DISE-inducing siRNAs into these cells. Knockdown of Ago2 did block this toxicity (**Fig. 4D**). It is possible that knockdown of Ago2 was insufficient to rescue the toxicity induced by constitutively expressed sequences. Thus, we tested Ago1, Ago2 and Ago 1/2/3 k.o. HCT116 cells to determine if cells completely devoid of these Argonaute proteins were resistant to the toxic effects of DISE-inducing si-or shRNAs. Knockout of Ago2 and triple k.o. of Ago 1/2/3 blocked the toxicity exerted by the siRNAs siL3 (2) and siGGGGGC [22], while Ago1 k.o. cells remained sensitive (**Fig. 4E**). However, Ago2 k.o. cells retained sensitivity to the DISE inducing shRNAs, shL1 and shL3 (**Fig. 4F**). It is possible that these shRNAs may exert toxicity through other Ago proteins. Interestingly, knockout of Ago 1/2/3 in HCT116 rescued shL1 toxicity but the cells remained somewhat sensitive to shL3 (**Fig. 4G**). This led us to wonder if the remaining Ago protein present in human cells, Ago4 may mediate DISE in HCT116 cells.

### CD95L kills HCT116 Ago1/2/3 k.o. cells

The association of the four Argonaute proteins in human cells with miRNAs has been found to be largely redundant [23, 24]. Due to this observation, the fact that Ago4 lacks endonuclease activity [25–27], and is often expressed at low levels in wt cells [24, 28], it is unclear if Ago4 mediates canonical RNAi or if it exerts unique cellular functions. Our data did present the possibility that Ago4 may be functional in mediating DISE in Ago 1/2/3 k.o. cells. We observed that Ago 1/2/3 k.o. cells were moderately sensitive to expression of pLenti-CD95L NP similar to what we saw for shL3 (**Fig. 5A**). Expression of the mRNA was similar between HCT116 wt and Ago 1/2/3 k.o. cells but increased in Drosha k.o. cells (**Fig. 5B**). To determine if CD95L could still be mediating toxicity through CD95L-derived sequences in the RISC, we performed Ago-RP-Seq. In the absence of Ago 1-3, Ago4 expression was substantially upregulated resulting in a dramatically increased pulldown of Ago4 in these k.o. cells (**Fig. 5C**). Upon analysis of the Ago4 bound sRNAs, we found very few CD95L-derived reads in the RISC (**Fig. 5D**), an observation seemingly inconsistent with RNAi being involved in the toxicity.

**Figure 5.**
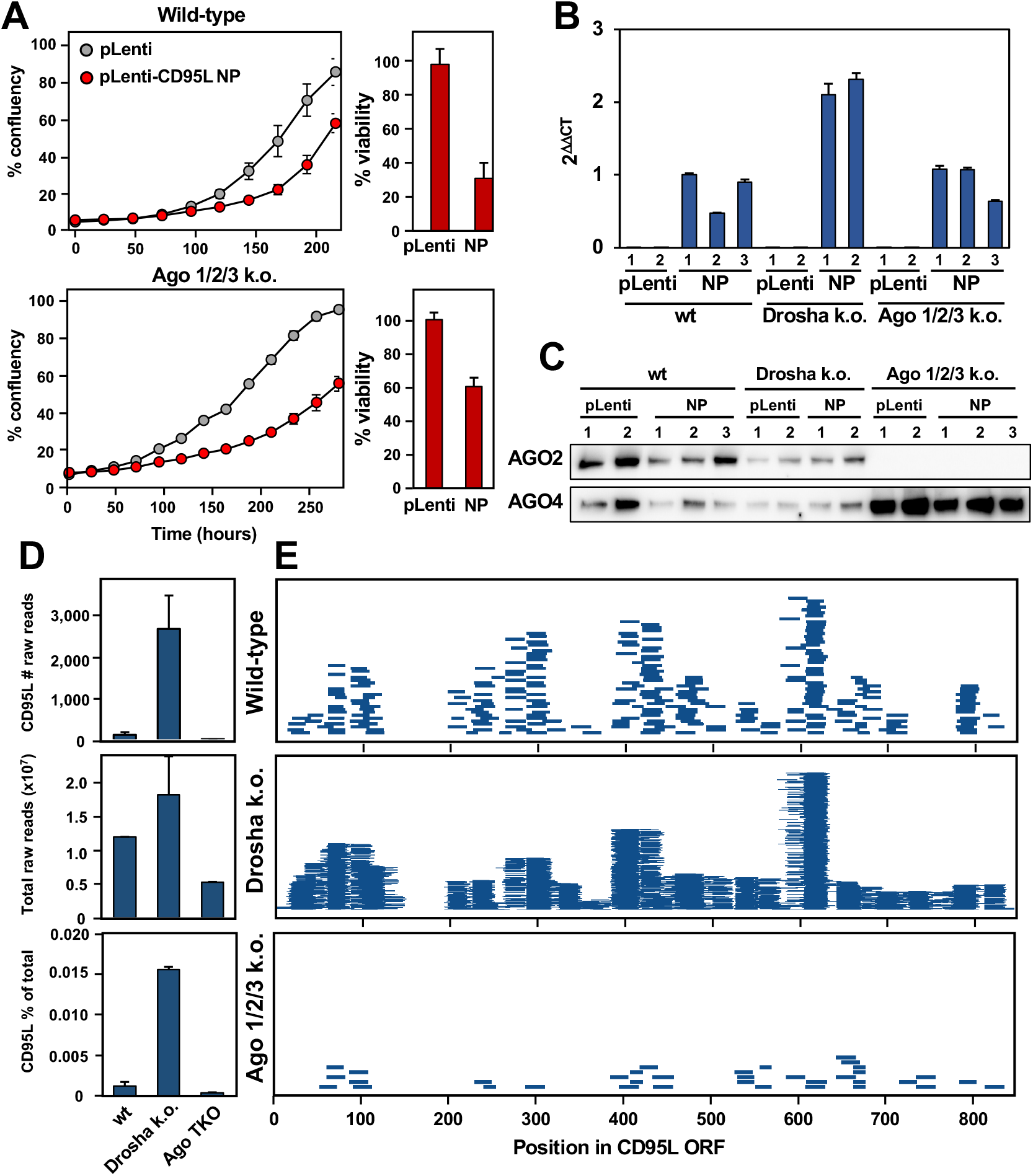
CD95L NP is toxic to HCT116 Ago 1/2/3 k.o. cells. **(A)***Left*, Percent confluency over time in HCT116 (top) and HCT116 Ago 1/2/3 k.o. cells (bottom) expressing pLenti empty vector control (pLenti) or pLenti CD95L NP (NP). *Right*, Relative cell viability at 144 hours. Error bars represent the standard deviation of triplicates. **(B)** Real-time qPCR analysis of CD95L expression in samples collected for Ago pulldown experiments. Bars represent the mean with standard deviation of triplicates. (**C)** Western blot analysis of Ago protein pulldown by the T6B peptide in HCT116, Drosha k.o., and Ago 1/2/3 k. o. cells. **(D)** Raw counts of R-sRNAs aligning to CD95L NP (top). The total raw R-sRNA reads sequenced per sample (center). The percentage of total raw R-sRNA reads that are derived from pLenti-CD95L NP (bottom). The average of two replicates are shown. Error bars represent the variance of the. **(E)** Mapping of CD95L NP-derived R-sRNAs along the CD95L ORF in the indicated genotypes. Each horizontal line represents one read. Reads from both replicates are displayed.

### CD95L-derived reads skew more toxic than reads derived from other mRNAs but the RISC favors nontoxic reads

While our data demonstrated that CD95L-derived sRNAs can exert toxicity in cells, it was unclear if this was a characteristic specific to CD95L mRNA [8, 17]. We previously reported that in HCT116 Drosha k.o. cells hundreds of protein coding genes are processed into sRNAs and loaded into the RISC [8]. Many of these genes are involved in protein translation and cell cycle control. To determine if the sRNAs derived from these protein coding genes are nontoxic, we applied our 6mer seed viability data to predict the effect their expression would have on cell fate. We identified the top 10 processed mRNAs by ranking coding genes by expression and processing [8]. As a way of measuring processing, we defined a stack as at least 10 raw reads mapping with the same 5’ start site to a gene. The top 10 most processed genes, in addition to CD95L, all had greater than 10 stacks per gene both in the RISC (**Fig. S7A**) and in the total sRNA (**Fig. S7B**). Applying a 6mer seed viability analysis to all sRNAs 18-25 nts long, we found that CD95L-derived reads skewed more toxic than those derived from the top 10 processed mRNAs (**Fig. S7C**). This was not merely a feature of exogenously expressed CD95L as endogenous CD95L-derived reads in pLenti control vector expressing cells were also similarly toxic (**Fig. S7C**, right panel) although present at much lower levels (**Fig. S7D**). A breakdown of the median 6mer seed viability of total cellular sRNAs by gene predicted that CD95L-derived sRNAs were more likely to exhibit toxicity through RNAi (**Fig. S7E**). However, it appeared that in the Drosha k.o. cells the most toxic sRNAs of the highly processed genes were less likely to be loaded into the RISC **(Fig. S7F)**. R-sRNAs derived from CD95L exhibited only a slightly lower median 6mer seed viability distribution than the ones derived from the top 10 genes (**Fig. S7F**, right panel). The median 6mer seed viability of CD95L-derived reads was 70%, while that of all Top 10 genes in aggregate was 73%. This contrasted to the median seed viability of all R-sRNAs which was 26%. We conclude that highly expressed processed protein coding genes are not favored to produce sRNAs that will exert toxicity through RNAi. Interestingly, while CD95L did produce sRNAs capable of exerting toxicity through RNAi, this analysis suggested that the majority of CD95L-derived R-sRNA were not likely to be toxic. This also suggested that CD95L mRNA expression may affect cell fate by changing the composition of R-sRNAs derived from other endogenous genes.

### Exploring the sequence determinants of CD95L toxicity

One of the main barriers to studying the toxic effects of CD95L mRNA is that the protein is a well characterized inducer of extrinsic apoptosis. To separate the effects of the mRNA from the protein, we previously generated various CD95L mutants. CD95L NP (**Fig. 6A**) exerts toxicity through a caspase-independent mechanism [8]. In the same study, we generated several additional CD95L mutants. CD95L Zero contained the same mutations as CD95L NP with an additional mutation in the first alternative start codon (AUG >AUA) (**Fig. 6A**). This blocked CD95L protein expression but retained toxicity [8]. We also hypermutated the CD95L sequence, introducing 303 synonymous mutations in CD95L SIL [8]. This mutant was still toxic. The only mutant we found to be non-toxic had all in-frame start codons mutated to stop codons. However, expression of this mutant was significantly reduced (data not shown). We suspected that the introduction of premature stop codons activated nonsense mediated decay (NMD) [29]. To avoid activating NMD, we mutated all in-frame start codons from AUG to GUA (**Fig. 6A**). Expression of this mutant, CD95L GUA produced mRNA at similar levels to wild-type CD95L (**Fig. 6B**), but CD95L GUA was nontoxic, and even appeared to promote cell growth (**Fig. 6D, E**).

**Figure 6.**
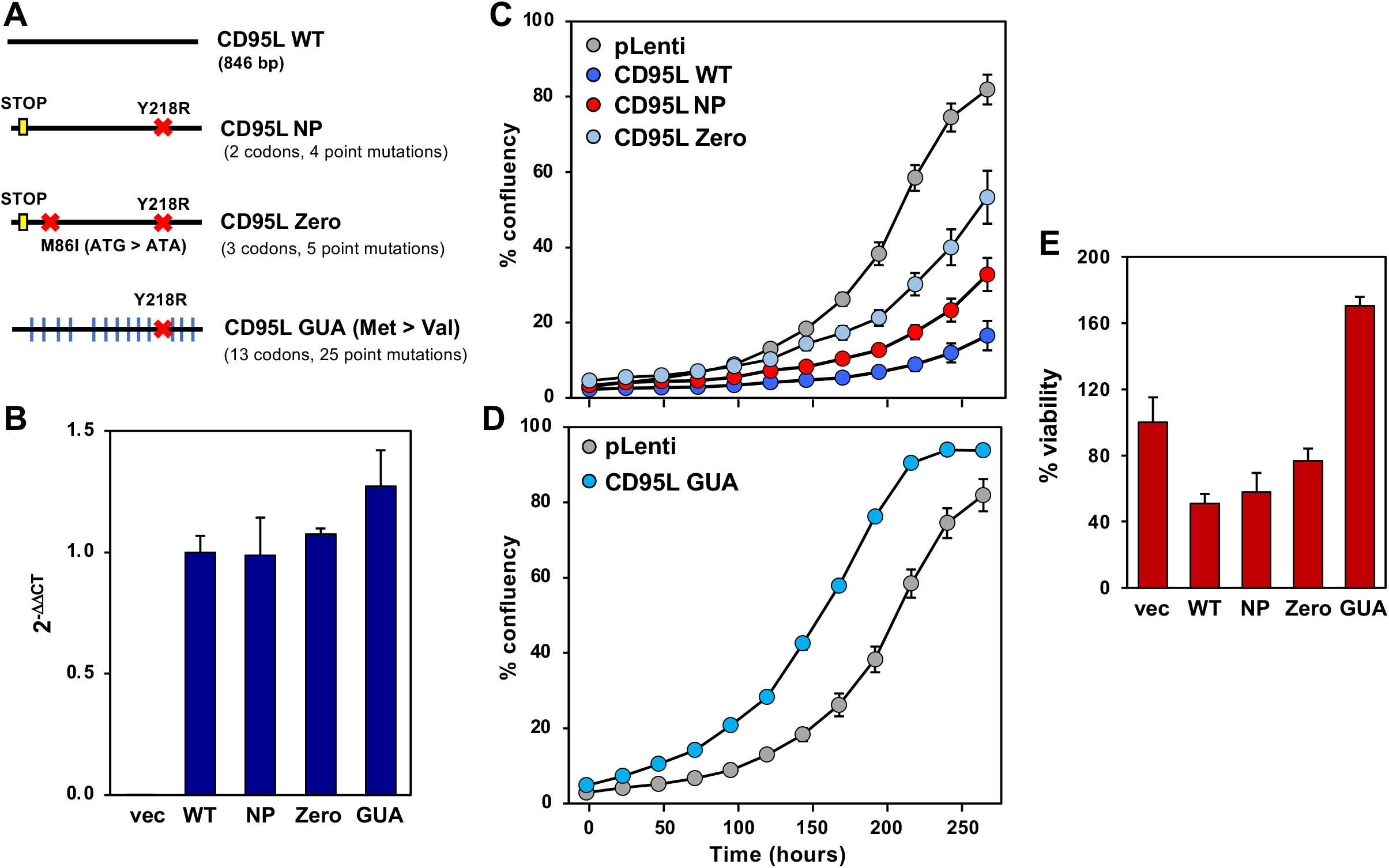
The mutant CD95L GUA is not toxic to cells. **(A)** Scheme representing the various CD95L mutants. A yellow box represents a mutation to a stop codon (UGA), an X indicates a point mutation, and a blue vertical line indicates mutation of an in-frame start codon (AUG) to valine (GUA). **(B)** Real-time qPCR analysis of CD95L mRNA expression in HCT116 Drosha CD95 d.k.o. c12 100 hours after infection with the CD95L mutants pLenti empty vector (vec), wild-type CD95L (WT), CD95L NP (NP), CD95L Zero (Zero), CD95L GUA (GUA). Expression was normalized to pLenti wild-type CD95L. Bars represent the standard deviation of triplicates. Data is representative of two independent experiments. **(C)** Percent confluency over time of HCT116 Drosha CD95 d.k.o. c12 infected with pLenti empty vector (vec), pLenti-CD95L WT, pLenti-CD95L NP, pLenti-CD95L Zero, and **(D)** pLenti-CD95L GUA. Error bars represent the standard error of triplicates. **(E)** Relative cell viability at 120 hours. Error bars represent the standard deviation of quadruplicates.

To determine if CD95L mRNA processing and loading into the RISC differed between the various mutants, we performed an Ago-RP-Seq analysis. The expression levels of CD95L-derived reads and their contribution to the total RISC-bound sRNA population was similar between mutants in the d.k.o. cells (**Fig. S8A**) and so was the average read length (**Fig. S8B**). Read stacks were also quite similar among the mutants with the exception of the SIL mutant (**Fig. S8C**). Stacks were often located close to stem regions (**Fig. S8C**, secondary prediction shown below each stack plot) suggesting that whatever is processing these mRNAs exhibits a preference for cleavage at double-stranded regions of the mRNA. All mutants showed a glaring lack of reads in the region spanning positions 133-225. In each case this region was enriched in cytosines (corresponding to the proline richness of the corresponding protein stretch) suggesting that C-rich regions either are not processed or not loaded into the RISC. These similarities between the mutants were also reflected in the average seed viability of all CD95L-derived reads in the RISC (**Fig. 7A)**. The seed viability associated with each CD95L mutant was quite similar, although the seed viability was slightly less toxic in the CD95L SIL mutant. Thus, indicating no correlation between the seed viability of R-sRNAs derived from CD95L and the functional effect of expression of each CD95L mutant on cell viability.

**Figure 7.**
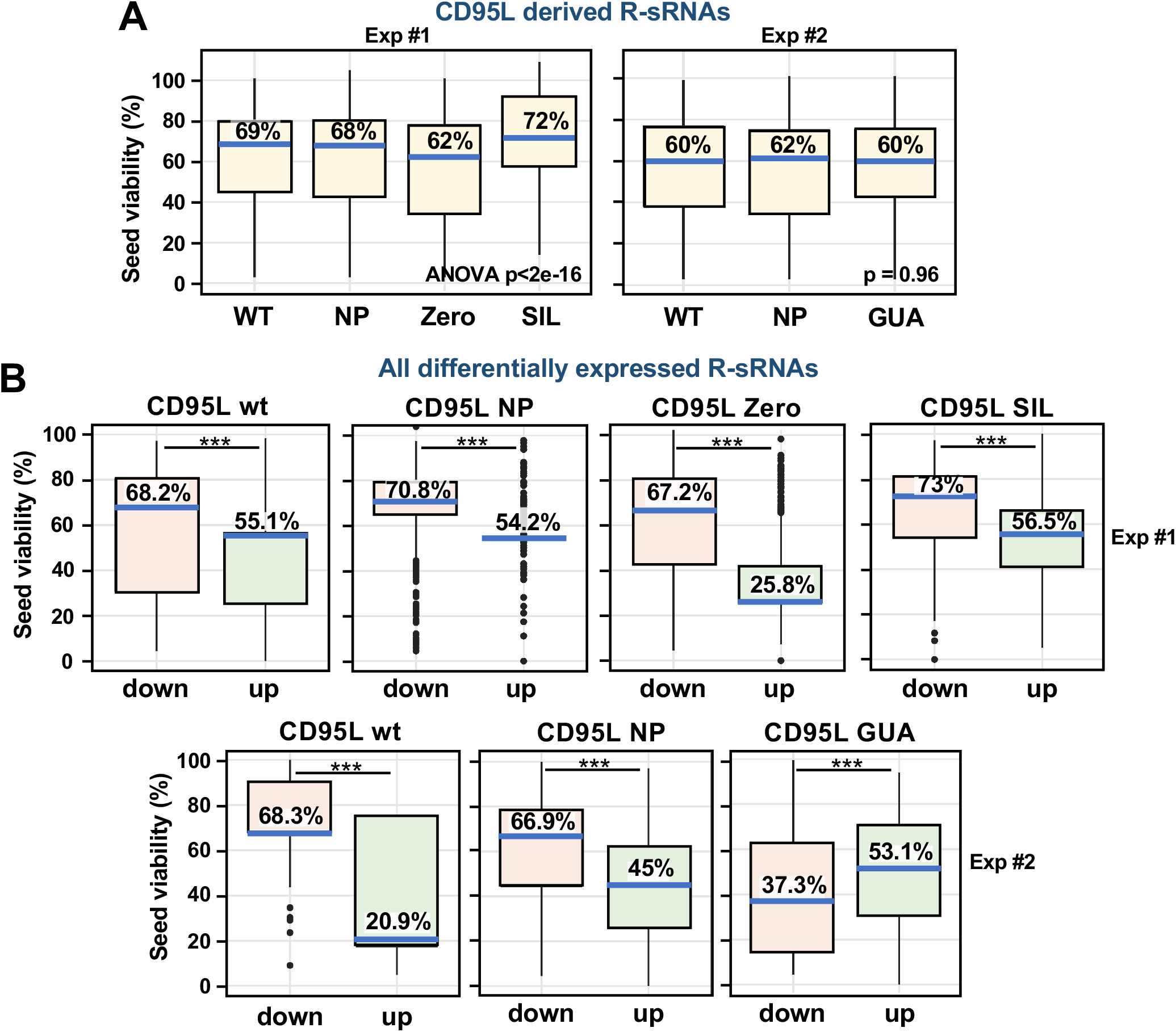
Shifts in the 6mer seed viability of R-sRNAs predict the toxic activity of mutant CD95L. **(A)** Boxplots with 6mer seed viabilities of the CD95L-derived sRNAs in the RISC. Reads from various CD95L mutants were compared. Data presents two independent experiments, experiment 1 (left) and experiment 2 (right). Kruskal-Wallis test *left*: p < 2 x 10^-16^ (the lower p-value threshold in R) and *right*: p = 0.951. **(B)** The 6mer seed viability of all differentially expressed R-sRNAs upon expression of various CD95L mutants. Only differentially expressed reads with an adjusted p-value < 0.05 are represented with the 6mer seed viability of down regulated reads in red and upregulated reads in green. Two independent experiments are represented with experiment one (top) and experiment two (bottom). Blue horizontal lines represent the median 6mer seed viability (% seed viability is given).

We recently provided evidence to suggest that in a number of models it is the balance of all RISC bound reads with toxic versus nontoxic seeds that can determine cell survival [14–16]. Using SPOROS we therefore analyzed the 6mer seed viability of reads that were significantly enriched or depleted (adjusted p-value < 0.05) in the RISC of d.k.o cells infected with the various CD95L mutants. In samples expressing toxic CD95L mutants, the median 6mer seed viability of the RISC shifted towards selective loading of more toxic sRNAs; Toxic R-sRNAs were significantly enriched and non-toxic R-sRNAs were downregulated (**Fig. 7B**). In contrast, the nontoxic CD95L GUA mutant, caused a shift towards loading of non-toxic sRNAs (**Fig. 7B**). These data suggest that a combination of CD95L-derived reads and reads derived from other genes determine cell fate upon introduction of CD95L mRNA into cells.

## Discussion

We previously reported that CD95L can kill cells independent of CD95L protein binding to its receptor CD95, and we provided evidence that the mRNA was killing cells through an RNAi-based mechanism similar to DISE [18]. We now report that CD95L mRNA processing results in production of the same shRNA sequence that led to the discovery of DISE [9]. This finding fueled further exploration of the role of the RNAi biogenesis machinery and Argonaute proteins in mediating CD95L mRNA processing and toxicity.

Our recent development of the Ago-RP-Seq-SPOROS pipeline [30] allows us to explore the connection between the sRNAs in the RISC and cell death in a standardized way. By applying data from the 6mer seed viability screens, we have now studied the connection between RISC bound sRNAs (R-sRNAs), their predicted effect on cell viability through seed-based targeting, and the resultant phenotypic outcomes in more detail. We reported that CD95L mRNA is processed into sRNAs with 6mer seed sequences that are predicted to be toxic to human cells [18]. We now find that CD95L-derived reads are more likely to exert toxicity than sRNAs derived from other highly expressed, processed, and RISC loaded endogenous mRNAs. We had experimentally determined that expression of CD95L-derived sRNAs as si-/shRNAs exert high levels of toxicity in cells. However, our data did not exclude that possibility that other endogenous R-sRNAs may contribute to this effect. We recently showed that shifts in the balance of R-sRNAs with toxic versus non-toxic 6mer seeds can effect phenotypic outcomes in therapy resistant ovarian cancer [14], in Alzheimer’s disease [15] and in HIV-1 infected cells [16]. Here we report that similar shifts in the balance of toxic versus non-toxic small RNAs in the RISC of CD95L expressing cells predict cell fate decisions. Expression of a number of toxic CD95L constructs resulted in shifts towards increased RISC loading of toxic sRNAs and decreased loading of non-toxic sRNAs. Conversely, the non-toxic CD95L mutant, CD95L GUA, resulted in a shift of R-sRNAs towards nontoxic sequences.

Based on previous observations in an ovarian cancer cell line [18] we assumed that CD95L mRNA toxicity was primarily mediated by Ago2, the only Argonaute protein with catalytic activity. Our observations reported here now suggest that this dependence on Ago2 may be cell type specific. Knockdown of Ago2 did not rescue CD95L toxicity in wt, Drosha k.o. of Drosha CD95 d.k.o. HCT116 cells. Neither knockdown nor knockout of Ago2 in HCT116 Drosha CD95 d.k.o. cells rescued toxicity exerted by DISE-inducing shRNAs. Only the activity of toxic DISE-inducing siRNAs was rescued by knockdown and knockout of Ago2. This may suggest that the kinetics or level of shRNA expression could overcome the loss of Ago2. Perhaps the persistent expression of an shRNA increases the likelihood the toxic shRNA exerts activity through catalytic deficient Ago proteins 1, 3, or 4. Because DISE results from seed-based targeting similar to the way that many miRNAs target genes, we might expect that catalytic-deficient Argonaute proteins would mediate DISE. Perhaps the kinetics of expression of certain shRNAs promotes loading of these DISE-inducing sRNAs by catalytic-deficient Ago proteins and the activation of DISE through seed-based targeting of survival genes.

Our data suggest that the HCT116 colon cancer cell line exhibits a special regulation of the RNAi pathway. The cells are uniquely amenable to genetic deletion of RNAi pathway components, which could be due to their ability to genetically compensate. For example, cells drastically upregulate Ago4 in response to the deletion of Ago 1-3 [16]. We hypothesize that this robustness may underlie the differential effects of Ago2 knockdown on CD95L toxicity. Work from other groups has suggested that Ago proteins could be functionally redundant [31]. It has been reported that miRNA binding does not differ much between Ago proteins [23]. The loss of Ago 1-3 in our report did however result in decreased loading of CD95L-derived reads into the RISC. Despite the reduction in CD95L reads cells remain sensitive to toxicity upon CD95L mRNA expression. We need to point out that while the number of CD95L derived R-sRNAs pulled down with Ago4 in the Ago 1,2,3 triple knock-out cells was very small, it was not smaller than the number of CD95L-derived R-sRNAs pulled down in CD95L infected HeyA8 cells which we established to die by through RNAi upon expression of CD95L NP [18]. Overall, our data suggest that death may result not only from loading of toxic CD95L-derived reads into the RISC, but also from increased loading of other sRNAs with toxic 6mer seed sequences.

A major unanswered question remains; what is processing mRNAs to produce R-sRNAs? The observation that CD95L-derived sRNAs map to regions of dsRNA in the predicted secondary structure of CD95L mRNA led us to hypothesize that Dicer could mediate processing. We performed Ago-RP-Seq in HCT116 Dicer k.o. cells expressing pLenti-CD95L NP and found no difference in the processing or loading of R-sRNAs. In fact, virtually the same CD95L-derived reads seen in the RISC of Drosha k.o. cells appeared to also be more abundant in the RISC of Dicer k.o. cells. Likewise, endogenous processed mRNAs, especially mRNAs involved in protein translation were also processed and loaded into the RISC in the absence of Dicer. Thus, the endonucleases that produce mature miRNAs, Drosha and Dicer, are not involved in the generation of R-sRNAs from protein coding mRNAs.

Multiple mechanisms of RNA degradation are coordinated at ribosomes such as nonsense-mediated decay, non-stop decay, and no-go decay [32]. These pathways degrade mRNAs that lack stop codons, contain premature stop codons, or transcripts with stalled ribosomes. Our observations involving the expression of various CD95L mutants suggest that CD95L processing and toxicity could involve regulation at ribosomes. In HCT116 cells we could not rescue toxicity by manipulating the RNAi pathway. Only changes to the sequence of CD95L prevented toxicity. It is interesting that the introduction of 303 synonymous point mutations had no effect on toxicity, although it did significantly change the pattern of mRNA processing. The only CD95L mutant that blocked toxicity was CD95L GUA. This mutant had all in-frame start codons mutated to valine codons. While these mutations did not affect the pattern of processing of this mutant, it is likely that these mutations would change the binding and translation of this transcript by ribosomes. Stalled ribosomes and ribosome collisions on mRNA transcripts have been shown to activate endonucleolytic cleavage of mRNA transcripts and could be involved in initiating the processing of CD95L mRNA and other mRNAs in the RISC. In fact, it has been demonstrated that phased cleavage products generated at stalled ribosomes persist in cells [33]. The endonuclease that mediates this processing, termed riboythrypsis, has yet to be identified. A possible candidate, the endonuclease NONU-1 (Cue2), has been shown to function in translational surveillance, and to cleave mRNA upstream of stalled ribosomes in no-go decay [34, 35]. However, loss of NONU-1 does not block the degradation of transcripts and data support a role for another yet to be identified endonuclease in this process [34]. Several RNA quality control pathways, such as No-Go decay and non-stop decay, have been shown to promote degradation of mRNA with stalled ribosomes. Mounting evidence suggests that several such pathways function redundantly, and that some pathways may only be activated in specific cell contexts, such during perturbation of another RNA quality control pathway [36], or in the absence of the Ribosome Quality Control Trigger complex [35]. Thus, identification of the endonuclease involved in the processed of R-sRNAs-derived from mRNAs may be difficult using loss of function genetic screens.

RNA degradation pathways are likely to be intricately connected to RNAi in cells. RNAi has been shown to promote mRNA transcript degradation by the decapping complex and the deadenylase complex [37, 38]. Increasing evidence supports a role for the non-stop translation surveillance pathway in the clearance of ribosomes from targeted transcripts and subsequent mRNA degradation [39–41]. Interestingly, this process appears to amplify the silencing activity of the RISC [42]. One could speculate a relationship may exist between the activation of RNA surveillance pathways and RNAi to prevent the translation of toxic proteins or peptides. Future work should determine the degree to which CD95L and other processed mRNAs are regulated by such RNA surveillance pathways. The link between known targets of no-go decay and RNAi could also be explored. If RNAi promotes more ribosome stalling and no-go decay, R-sRNAs derived from mRNAs may further promote RNAi, or they could function as inhibitors of canonical RNAi. The idea that mRNA degradation products are not merely recycled but could in turn regulate gene expression is intriguing and could contribute to our understanding of the role of RNA in the regulation of gene expression at homeostasis and in the context of disease.

## Materials and methods

### Reagents and antibodies

Reagents used were puromycin (Sigma, #P9620), doxycycline (Sigma, #9891), leucine-zipper tagged (Lz)CD95L described [43], Lipofectamine RNAiMAX (ThermoFisher Scientific, #13778150), Lipofectamine 2000 (ThermoFisher Scientific, #11668027), polybrene (Sigma, #H9268). The following antibodies were used: PE conjugated-anti-CD95 (eBioscience, #12-0951-83; RRID: AB_465789); PE conjugated Isotype control (eBioscience, #12-4714-82; RRID: AB_470060); anti-β-actin antibody (Santa Cruz, #sc-47778, RRID: AB_626632); anti-AGO2 (Abcam, #32381 AB_867543), anti-AGO4 (Cell signaling #6913, RRID: AB_10828811), Goat anti-rabbit IgG-HRP (Southern Biotech, #SB-4030–05).

### Cell culture

293T cells were cultured in DMEM medium (Corning #10-013-CM) with 10% Serum Plus II Medium Supplement (Sigma-Aldrich #14009C). The following HCT116 cells were purchased from the Korean Collection for Type Cultures (KCTC): HCT116 wild-type (KCTC, cat #HC19023), HCT116 Drosha k.o. clone #40 (KCTC, cat #HC19020), HCT116 Dicer k.o. clone #43 (KCTC, cat #HC19023). The HCT116 Ago2 k.o. cells [44] were a kind gift from Joshua T. Mendell (UT Southwestern). The HCT116 Ago1 k.o. and HCT116 Ago 1/2/3 k.o. cells [28] were provided by David Corey (UT Southwestern). All HCT116 cells and knock-out clones were cultured in McCoy’s 5A medium (ATCC #30-2007) with 10% Serum Plus II Medium Supplement (Sigma-Aldrich #14009C). Cells were cultured at 37°C, 5% CO_2_ were tested for mycoplasma using PlasmoTest (Invitrogen). Before use the Serum Plus II Medium Supplement was heat inactivated by incubation at 56°C for 30 minutes.

### CRISPR-Cas9 deletion of CD95

HCT116 Drosha CD95 d.k.o. pool was generated by Synthego using the guide RNA sequence: GGAGUUGAUGUCAGUCACUU. Single cells were sorted into 96-well plates with 50% conditioned media by FACS. Colonies were treated with LzCD95L for 24 hours, and visually inspected for signs of apoptosis. Resistant clones were expanded and assayed for DNA editing by PCR. The following PCR primers were used to confirm deletion, forward: tggtgctgtttctagtgTGGt, reverse: TGTTGCTACTCCTAACTGTGACT (design: Synthego, Synthesis: IDTDNA, custom DNA oligos with standard desalting). ChoiceTaq Blue (Denville cat #C775Y30) master mix was used for PCR amplification following manufacturer’s protocol. The QIAquick PCR Purification Kit (Qiagen cat. #28106) was used to isolate the PCR product following the manufacturer’s protocol. The forward primer was submitted with the PCR product for Sanger sequencing (ACGT Inc.) The Synthego ICE CRISPR Analysis tool was used to analyze the editing efficiency (Synthego).

### CD95L lentiviral vectors

Generation of pLenti CD95L, pLenti CD95L NP, pLenti CD95L SIL, and pLenti CD95L Zero was described in [18]. All CD95L sequences were sub-cloned into the pLenti-GIII-CMV-RFP-2A-Puro vector (ABM Inc.). CD95L GUA was synthesized with NheI and XhoI flanking restriction enzyme sites and sub-cloned (GenScript). CD95L GUA contains the Y218R mutation (TAT > CGT) [1] and 13 in-frame Met > Val mutations (AUG > GUA).

### Lentivirus transduction experiments

Lentiviruses were packaged in 293T cells using psPAX2 (Addgene #12260) and pMD2.G (Addgene #12259) packaging plasmids. 293T cells were seeded to 90% confluency in 10 cm cell culture treated plates (Greiner Bio-ONE, cat #664160). Plasmids were introduced by transfection with 60 μl Lipofectamine 2000 (ThermoFisher Scientific, #11668027) diluted in 2 ml Opti-MEM (Gibco cat, # 31985-070) and 6 μg psPAX2, 3 μg pMD2.G packaging vectors, plus 10 μg delivery vectors, pLenti (ABM Inc.) or pTIP.[45]. Media was refreshed 18 hours later, and virus supernatant was collected after another 24 and 48 hours. Viral supernatant was centrifuged and filtered through a 0.45 μm PVDF membrane (MilliporeSigma, SLHVM33RS) to remove debris, and stored at - 80°C. Mission shRNA Lentiviral particles designed to target CD95L were purchased: shL3 (Sigma TRCN0000059000), shL1 (Sigma TRCN0000058999), and shScr (Sigma SHC002V). All transductions were performed in the presence of 8 μg/ml polybrene. HCT116 cells were resuspended in media with polybrene with 150,000 cells seeded in 24-well plates, 300,000 in 12-well or 500,000 cells seeded in 6-well plates. Virus was added by volume with either 25% or 50% virus optimized by viral titer. For commercial viruses the volume required to achieve a multiplicity of infection (MOI) of 2-3 was added to each well. Cells were incubated at 37°C, 5% CO_2_ overnight, and the viral media was replaced with regular media. Cells were treated with 2 μg/ml puromycin when RFP reporter expression was visible, usually 48-72 hours post infection. Cells were selected for 48 hours. After selection cells were reseeded for assays (time point 0).

### Transfection with short oligonucleotides

Custom siRNA oligonucleotides were ordered from integrated DNA technologies (IDT) as described previously: siCD95L cluster 15.1 and 15.2 [18], siL3 [9], siGGGGGC [12]. The non-targeting control siNT8 anti-sense 5’-UAAUCUAACAUGUAAACCAAA-3’, sense 5’-mUmGGUUUACAUGUUAGAUUATT-3’. A lowercase m before the base indicates 2’-O-methylated nucleotides. The oligos were annealed according to the manufacturer’s instructions. Duplexed siRNAs were introduced via reverse transfection using RNAiMax diluted in Opti-MEM in 96-well plates. A 50 μl transfection mix was added per well with a final siRNA concentration of 10 nM. The optimal volume of RNAiMax was determined for each clone, 0.2 or 0.3 μl was used. Diluted cells (3,000 – 8,000) were plate in 200 μl in each well on top of the transfection mix to a final volume of 250 μl. The effect on cell growth and viability was assayed as described below.

### Ago2 knockdown experiments

HCT116 wt and Drosha k.o. cells were reverse transfected with 25 nM siRNA SMARTpool targeting siAgo2 (Dharmacon, L-004639-00-0005) or a control siRNA pool (siCtr, Dharmacon, D-001810–10). The transfection mix was prepared in 500 μl Opti-MEM with 2 μl RNAiMax. Cells were seeded 500,000 cells in 1.5 ml media. The cells and transfection mix were combined in 6-well plates. Cells were incubated at 37°C, 5% CO_2_ overnight, and the media was replaced the following day. The following day (48 hours post-transfection) cells were transduced with 25% pLenti of pLenti CD95L NP, cells were seeded in 12-well plates with 250,000 cells per well in 750 μl media, 8 μg/ml polybrene and 250 μl of virus was added. After an incubation at 37°C, 5% CO_2_ overnight the viral media was replaced. Cells were treated with 2 μg/ml puromycin for 48 hours and then plated for IncuCyte Zoom analysis of cell growth. HCT116 Drosha CD95 d.k.o. clone #12 were prepared with the following modifications. To knockdown Ago2, cells were reverse transfected in 24-well plates, 150,000 cells were plated per well. The SMARTpool siRNAs were diluted to 20-25 nM per well, and cells were transduced with 50% pLenti or pLenti CD95L NP viruses. For shRNA transductions, cells were infected with pLKO-shScr or shL3 at an MOI of 3.0. Uninfected cells were harvested 96 hours after transfection to assess the efficiency of Ago2 knockdown. Cell viability was determined 120 hours after puromycin selection.

### Analysis of cell growth and viability

Cells were seeded in 96-well plates, 2000 to 8000 cells per well, in triplicates or quadruplicates. For analysis by IncuCyte cells were seeded in TPP tissue culture test plates (TPP, cat #92696). For analyses of ATP content cells were plated in either white CELLSTAR tissue culture microplates (Greiner, BIO-ONE, cat # 655083) or μCLEAR Black, CELLSTAR, tissue culture microplates (Greiner, BIO-ONE, cat # 655090). Cell growth over time was analyzed using the IncuCyte ZOOM live-cell imaging system (Essen BioScience). Images were acquired every 4-6 hours using a 10x objective. The IncuCyte ZOOM software (version 2015A) was used to process images. Growth was analyzed as a change in cell confluency over time.

Cell viability was determined by the using CellTiter-Glo 2.0 Cell Viability Assay according to the manufacturer’s instructions (Promega, cat # G9241). Unless otherwise indicated, ATP content was assayed at 96 hours after seeding. Briefly, media was removed to reduce the cell culture media to 75 μl per well and 75 μl of the CellTiter-Glo reagent was added. The plate was agitated on an orbital shaker for 2 minutes and luminescence was read after a 15-minute incubation at room temperature using the BioTek Synergy Neo2.

### Real-time PCR

Real-time PCR was performed to verify CD95L expression, and to compare the relative expression of the various CD95L mutants. To make cDNA, the High-Capacity cDNA Reverse Transcription Kit (Applied Biosystems #4368814) was used with 200 ng input total RNA, determined by nanodrop readings (Nanodrop 2000). The qPCR reaction was performed using Taqman gene expression mix (ThermoFisher Scientific, #4369016) and the following FAM Taqman probes: GAPDH (Hs00266705_g1), human CD95L (Hs00181226_g1 and Hs00181225_m1), and a custom probe to detect the CD95L SIL (Thermofisher Scientific, assay ID: APNKTUD [18]). Reactions were performed in triplicate in 96-well plates (Applied Biosystems, cat # N8010560) in the Applied Biosystems 7500 Real Time PCR system. CT values were determined using default settings, and relative expression was determined using the ΔΔCT method normalized to samples expressing pLenti CD95L wild type.

### Western Blotting

To confirm Ago2 knockdown, cells were lysed in Killer RIPA lysis buffer (150 mM NaCl, 10 mM Tris-HCl pH 2.7, 1% SDS, 1% Triton X-100, 1% deoxycholate, 5 mM EDTA) with freshly added protease inhibitor cocktail (Roche, cat #11836170001). Lysates were passed through a 26G syringe and protein was quantified by DC Protein Assay (Bio-Rad, cat #5000112). Lysates were diluted to equal volumes and 30 ug protein was loaded per well using the Novex WedgeWell Tris-Glycine Welcome Pack (ThermoFisher Scientific, cat #XP0420B). The SDS-PAGE gel was transferred to a nitrocellulose membrane (Sigma, cat #GE10600016). These blots were blocked in 5% milk TBS-T (0.1% Tween-20/TBS) and incubated overnight at 4°C in primary antibodies diluted in 5% milk TBS-T: anti-Ago2 (1:500-1000), anti-Ago4 (1:1000), anti-Actin (1:5000). The blots were incubated in the secondary antibody for 2 hours at room temperature, anti-rabbit IgG HRP diluted (1:5000) in 5% milk TBS-T. Protein bands were visualized using the SuperSignal West Dura Extended Duration Substrate (ThermoFisher Scientific, cat #34076) captured by the G:Box Chemi XT^4^ imager (Syngene).

### CD95 Surface Staining

Cells were resuspended in 300 μl 5% BSA in PBS with 5 μl of anti-CD95 or IgG control. Cells were incubated for ~4 hours on ice. Then cells were fixed in 4% formaldehyde in PBS for 20 minutes. Cells were washed three times with PBS and resuspended in 5% BSA in PBS. Cells were kept on ice and analyzed on the BD LSRFortessa SORP Cell Analyzer with HTS.

### Sample preparation for Ago-RP-Seq

Spin infections were performed with cells in suspension in 6-well plates. HCT116 cells (wt, Drosha k.o., Dicer k.o. and Drosha/ CD95 d.k.o. clone #12) were passed through a 40 μm cell strainer and diluted to 500,000 cells per ml in media with 16 μg/ml polybrene. Cells were plated with 1 ml virus or DMEM media control (final concentration of polybrene 8 μg/ml) in 6-well plates with 20-36 wells infected per virus. The plates were centrifuged at 2700 rpm for 20 minutes at room temperature, and incubated at 37°C, 5% CO_2_ overnight. The next day cells were trypsinized and pooled, 10-18 wells per sample replicate (two sample replicates per virus) and replated in 15 cm dishes. Cells were treated with 2 μg/ml puromycin 48-72 hours after infection. Cells were expanded as needed. After 48 hours of puromycin selection cells were plated for IncuCyte analysis and ATP assay. On day 5 (~122 hours post-infection), cell pellets were harvested. Cells were trypsinized, pelleted, washed with PBS and counted. Aliquots of 8-10 million cells were pelleted and flash frozen for Ago-RP. Note: Experiments with more than two sample replicates (replicates 3-4) are biological replicates.

### Ago-RP and library preparation for small RNA-Seq

Pulldown of Ago 1-4 was performed as previously described [18]. Briefly, cell pellets were lysed in 1 ml NP-40 lysis buffer (50 mM Tris pH 7.5, 150 mM NaCl, 5 mM EDTA, 0.5% NP-40, 10% (v/v) glycerol, 1 mM NaF; supplemented with 1:200 EDTA-free protease inhibitors (Millipore, #539134) and 1:1000 RNaisin Plus (Promega, #N2615). Samples were vortexed and kept on ice ~30 minutes. Cell debris was removed by centrifugation at 20,000 x G for 20 min. at 4°C. Lysates were transferred to Lo-bind tubes (Eppendorf, #022431021) and incubated with 500 μg Flag-GST-T6B peptide [46] and 80 μl of anti-Flag M2 Magnetic beads (Sigma, #M8823) for at least 3 hours rotating at 4°C. Beads were washed three times with 1 ml NP-40 lysis buffer. After the last wash, 100 μl were aliquoted for western blot analysis to confirm the efficiency of Argonaute pulldown. The NP-40 lysis buffer was removed and beads were resuspended in 500 μl Trizol (Ambion, #15596018). RNA was extracted according to the manufacturer’s protocol, resuspended in 20 μl UltraPure water and divided into two 10 μl aliquots.

Size markers 19 and 35 nucleotides long were dephosphorylated using 0.5 U/ μl CIP alkaline phosphatase (NEB M0290L) by incubating at 37°C for 15 minutes, and then radiolabeled with 0.5 μCi (γ-^32^P)ATP (Perkin Elmer NEG002Z250UC) using T4 PNK kinase (NEB M0201L) incubating at 37°C for 20 minutes. The size markers were resolved on a 15% urea-PAGE gel (National Diagnostics, EC830, EC835, EC840), extracted, resuspended in UltraPure water for use in subsequent steps. The RNA isolated from the Ago-RP was then prepared for small RNA sequencing as previously described [47]. Briefly, the RNA and the size marker was ligated with Barcoded 3’ adenylated adapters using the T4 RNA Ligase 2, truncated K227Q (NEB, #M0351) for at least 3 hours at 16°C. The ligation products were pooled into one tube and isolated by ethanol precipitation. The RNA-adapter pellet was dissolved in 20 μl water and resolved on a 15% urea-PAGE gel flanked by radiolabeled size markers with adapters. RNA the size of the shifted size markers was excised and a gel extraction was performed. The RNA-adapter pellet was resuspended in 9 μl UltraPure water. A 5’ adapter was ligated using T4 RNA ligase, T4 Rnl1 (Thermofisher, cat #EL0021) incubated at 37°C for 1 hour while agitating. The product was resolved on a 12% urea-PAGE gel flanked by 3’ and 5’ ligated radiolabeled size markers. The small RNA adapter product was excised and a gel extraction was performed. The RNA-adapter pellet was resuspended in 4.6 μl UltraPure water, and reverse transcribed using the Superscript III reverse transcriptase (Invitrogen, cat #18080-044). The cDNA was amplified by PCR using Platinum^™^ *Taq* DNA Polymerase (Invitrogen, cat #10966-018). Single end sequencing was performed on the Illumina Hi-Seq 4000.

### Processing of small RNA-Seq data

The raw small RNA-seq data was processed as previously described [18]. Illumina adapter sequences were removed with trim_galore. Unique Molecular Identifiers (UMIs) four nucleotides 5’ of the sequenced small RNA and two nucleotides 3’ of the sequence were removed. Tophat was used to align reads to the hg38 assembly of the human genome. Raw read counts were assigned to genes using HTseq. Gene expression was normalized based on library size and sequence complexity using EdgeR.

### Identification of CD95L reads

CD95L reads were identified as previously described [18]. Briefly, the reads from each sample were compiled as a BLAST database, and blastn was utilized to match the CD95L ORF (wild type NM_00639.2 or CD95L mutant sequences) to reads in each sample. Reads were considered matches if they had an e-value < 0.05% and 100% identity across the length of the read. These hits were converted to bed files that were used for subsequent analyses. Sample replicates were combined and stack plots depicting the locations where RISC bound CD95L sRNAs map along the CD95L ORF were generated using the R package Sushi.

### Analysis of CD95L read abundance

CD95L read abundance was determined by tallying raw matching reads identified using blastn. CD95L reads were counted for each sample from individual replicate bed files. The total number of raw reads per sample was calculated, as was the percent contribution of CD95L reads to the total read counts.

### Identification of shL3 reads

The presence of CD95L-derived sRNAs with the same sequence as commercial CD95L targeting shRNAs was queried by searching the bed files for reads that contain the 5-p arm of the shRNA. Reads mapping to cluster 15 in the CD95L ORF contained the shL3-5p sequence (with the same ‘5 start site, −3,+1) were tallied and were highlighted in orange in Fig. 1A using Sushi. The percentage of reads mapping to shL3 from CD95L was calculated, and the variation in the 5’ end of mapping reads was depicted in the pie chart in Fig. 1B.

### CD95L mRNA processing analysis

Analysis of small RNA-Seq data of shL3 infected cells was described previously [9]. CD95L read length was determined from sample .bed files and tallied using the tidyverse package in R. For the analysis of the start and stop site of abundant stacks in HCYT116 wt, Drosha k.o., and Dicer k.o. cells the top 10 most abundant stacks were identified. The most abundant read was annotated as starting at position 0 based on the mapping of the 5’ end of the read. Using the mapping of the 5’ start site indicated in sample .bed files the number of reads mapping −5 to +5 nucleotides was tallied, and plotted as line plots using ggplot2 in R. To analyze processing at the 3’ end of each read, reads starting at position 0 were analyzed. The read length of each was tallied and also plotted as line plots.

### CD95L stem-loop and nucleotide analysis

RNAfold from the ViennaRNA package was used to predict the secondary structure of CD95L mRNA and the various CD95L mutants [48]. The RNAfold Dot-Bracket output was imported into R. A barplot of height equal to 1 (y-axis) was generated using the package ggplot2 with one bar representing each nucleotide in the CD95L ORF (846 in total on the x-axis). The color of the bars correspond to the Dot-Bracket notation, blue indicates unpaired nucleotides (dots), and red bars indicate paired nucleotides (brackets). The same strategy was employed to visualize the nucleotide composition of the CD95L sequences. Again, ggplot2 was used to generate a bar plot with a x-axis 1-846 and a y-axis equal to 1. The color of the bars represent different nucleotides bases in the CD95L sequence with **A**denines in blue, **C**ytosines in yellow, **G**uanines in red, and **U**racils in green.

### CD95L 6mer seed toxicity analysis

The six nucleotide seed sequence (positions 2-7) was extracted from the reads in the CD95L bed files. The 6mer seed viability data corresponding to each 6mer seed sequence from the average of three human cell lines (6merdb.org) was added in R using the tidyverse package. The viability data was summarized in boxplots using ggplot2. Only reads 18-25 nucleotides long were included in the analysis. A breakdown of the contribution of different seed sequences to the overall seed viability was profiled by binning the viability data and counting the number of reads expected to exert the same effect on cell viability. The viability data was rounded to the nearest integer, and the number of reads with the same viability was tallied. The binned data was then plotted in Excel with viability on the x-axis and read counts on the Y-axis as described in [13].

### Identification of processed mRNAs

For each sample, small RNA-Seq reads were aligned to the human genome build hg38 using Tophat and converted to .bam files. Using blastn, each .bam file was compiled into a blast database. A .fasta file containing the longest annotated transcript of each protein coding gene was queried against each sample database. Matches were filtered for 100% identity and an e-value < 0.05%. Only uniquely mapping reads were considered hits. Regions with greater than 10 reads mapping with the same 5’ start site were defined as a stack. The number of stacks, or the stack count, across a gene transcript was tallied, and the stack counts were merged with the normalized RNA-Seq tables. The table was annotated with gene biotypes information from Ensembl using biomart. A processed mRNA was identified as any gene annotated as protein-coding with three or more stacks and 10 or more normalized reads in averaged sample replicates. Scatterplots representing stacks by normalized gene expression were generated in R using ggplot2. Dark blue data points represent genes that were common in HCT116 wt, Drosha k.o. and Dicer k.o. cells. A Venn analysis was performed using the web-based application BioVenn [49]. The Database for Annotation, Visualization and integrated Discovery (DAVID) release v2022q2 was utilized to analyze the processed gene list from each genotype for enriched gene ontologies [50, 51]. Gene ontologies were annotated using the GOTERM_BP_FAT. Common GOterms with greater than 1.5 fold enrichment and a Bonferroni adjusted p-value < 0.05 were considered to be significantly enriched. The top 10 processed genes described in Fig. S7 were identified by ranking processed mRNAs by stack count and expression. Genes ranked in the top 50 in both the Ago pulldown and total small RNA-Seq data were selected for further curation. Genes with reads annotated in miRBase (v. 22.1) as miRNAs were excluded. Also genes with reads mapping primarily to one location were excluded.

### Analysis of reads from processed mRNAs

As described for CD95L, BLAST databases compiled for each sample replicate were queried for matches to select protein-coding mRNA transcripts. Hits with an e-value < 0.05% and 100% identity across the read were compiled in .bed files. .bed files were concatenated and plotted using the Sushi package in R. For analysis of the predicted effect of mRNA reads on cell fate, the six nucleotide seed sequence (positions 2-7) was extracted from each read and the 6mer seed viability data, averaged across human samples, corresponding to each seed sequence was matched. A data frame was generated with the 6mer seed viability “score” printed one time for each read in a sample. The data was represented as density plots, boxplots or violin plots using the R package ggplot2.

### Statistical analyses

Student’s T-tests were performed in Microsoft Excel. All other statistical analyses were conducted in R version 4.0.0.

## Data availability

## Acknowledgements

We grateful for help from members of this core facility, Pankaj Bhalla, Ph.D. and Alex Yemelyanov, Ph.D. Thanks also to Sergii Pshenychnyi, Ph.D. and the Northwestern University—Recombinant Protein Production Core for help with optimizing production of the GST-T6B-flag peptide. This work was supported by the Northwestern University – Flow Cytometry Core Facility supported by Cancer Center Support Grant (NCI CA060553). Flow Cytometry Cell Sorting was performed on a BD FACSAria SORP system and BD FACSymphony S6 SORP system, purchased through the support of NIH 1S10OD011996-01 and 1S10OD026814-01. Research reported in this publication was supported by Northwestern University Skin Biology & Diseases Resource-Based Center of the National Institutes of Health under award number P30AR075049.

## Author’s Contributions

Material preparation, data collection and analysis was performed by Ashley Haluck-Kangas and Madelaine Fink; bioinformatic analyses were designed by Elizabeth Bartom and performed by Ashley Haluck-Kangas; the Manuscript was drafted, revised and edited by Ashley Haluck-Kangas and Marcus Peter; the study was conceptualized by Marcus Peter.

## References

1. Schneider P, Bodmer JL, Holler N, Mattmann C, Scuderi P, Terskikh A, Peitsch MC and Tschopp J (1997) Characterization of Fas (Apo-1, CD95)-Fas ligand interaction. J Biol Chem 272:18827–33.

2. Algeciras-Schimnich A, Shen L, Barnhart BC, Murmann AE, Burkhardt JK and Peter ME (2002) Molecular ordering of the initial signaling events of CD95. Mol Cell Biol 22:207–20.

3. Nagata S (1997) Apoptosis by death factor. Cell 88:355–65.

4. Fulda S, Scaffidi C, Pietsch T, Krammer PH, Peter ME and Debatin KM (1998) Activation of the CD95 (APO-1/Fas) pathway in drug-and gamma-irradiation-induced apoptosis of brain tumor cells. Cell Death Differ 5:884–93.

5. Muller M, Strand S, Hug H, Heinemann EM, Walczak H, Hofmann WJ, Stremmel W, Krammer PH and Galle PR (1997) Drug-induced apoptosis in hepatoma cells is mediated by the CD95 (APO-1/Fas) receptor/ligand system and involves activation of wild-type p53. J Clin Invest 99:403–13.

6. Kischkel FC, Hellbardt S, Behrmann I, Germer M, Pawlita M, Krammer PH and Peter ME (1995) Cytotoxicity-dependent APO-1 (Fas/CD95)-associated proteins form a death-inducing signaling complex (DISC) with the receptor. EMBO J 14:5579–88.

7. Muzio M, Chinnaiyan AM, Kischkel FC, O’Rourke K, Shevchenko A, Ni J, Scaffidi C, Bretz JD, Zhang M, Gentz R, Mann M, Krammer PH, Peter ME and Dixit VM (1996) FLICE, a novel FADD-homologous ICE/CED-3-like protease, is recruited to the CD95 (Fas/APO-1) death-inducing signaling complex. Cell 85:817–27, shared senior authorship.

8. Putzbach W, Haluck-Kangas A, Gao QQ, Sarshad AA, Bartom ET, Stults A, Qadir AS, Hafner M and Peter ME (2018) CD95/Fas ligand mRNA is toxic to cells. Elife 7. doi: 10.7554/eLife.38621

9. Putzbach W, Gao QQ, Patel M, van Dongen S, Haluck-Kangas A, Sarshad AA, Bartom E, Kim KY, Scholtens DM, Hafner M, Zhao JC, Murmann AE and Peter ME (2017) Many si/shRNAs can kill cancer cells by targeting multiple survival genes through an off-target mechanism. eLife 6: e29702.

10. Hadji A, Ceppi P, Murmann AE, Brockway S, Pattanayak A, Bhinder B, Hau A, De Chant S, Parimi V, Kolesza P, Richards JS, Chandel N, Djaballah H and Peter ME (2014) Death induced by CD95 or CD95 ligand elimination. Cell Rep 10:208–222.

11. Gao QQ, Putzbach WE, Murmann AE, Chen S, Sarshad AA, Peter JM, Bartom ET, Hafner M and Peter ME (2018) 6mer seed toxicity in tumor suppressive microRNAs. Nat Commun 9:4504. doi: 10.1038/s41467-018-06526-1

12. Patel M, Bartom ET, Paudel B, Kocherginsky M, O’’Shea KL, Murmann AE and Peter ME (2022) Identification of the toxic 6mer seed consensus in human cancer cells. Sci Rep 12:5130.

13. Bartom ET, Kocherginsky M, Paudel B, Vaidyanathan A, Haluck-Kangas A, Patel M, O’Shea KL, Murmann AE and Peter ME (2022) SPOROS: A pipeline to analyze DISE/6mer seed toxicity. PLoS Comput Biol 18:e1010022. doi: 10.1371/journal.pcbi.1010022

14. Patel M, Wang Y, Bartom ET, Dhir R, Nephew KP, Adli M, Matei D, Murmann AE, Lengyel E and Peter ME (2021) The ratio of toxic-to-nontoxic microRNAs predicts platinum sensitivity in ovarian cancer. Cancer Research 81:3985–4000.

15. Paudel B, Jeong SY, Pena Martinez C, Rickman A, Haluck-Kangas A, T. BE, Frederiksen K, Affaneh A, Kessler JA, Mazzulli JR, Murmann AE, Rogalski E, Geula C, Ferreira A, Heckmann BL, Green DR, Sadleir KR, Vassar R and Peter ME (2022) Death induced by survival gene elimination (DISE) contributes to neurotoxicity in Alzheimer’s disease. BioRxiv at https://www.biorxiv.org/content/10.1101/2022.09.08.507157v1.

16. Vaidyanathan A, Taylor HE, Hope TJ, D’Aquilla RT, Bartom ET, Hultquist JF and Peter ME (2022) Contribution of 6mer seed toxicity to HIV-1 induced cytopacitity. BioRxiv at https://biorxiv.org/cgi/content/short/2022.10.01.510471v1.

17. Putzbach W, Gao QQ, Patel M, van Dongen S, Haluck-Kangas A, Sarshad AA, Bartom ET, Kim KA, Scholtens DM, Hafner M, Zhao JC, Murmann AE and Peter ME (2017) Many si/shRNAs can kill cancer cells by targeting multiple survival genes through an off-target mechanism. Elife 6. doi: 10.7554/eLife.29702

18. Putzbach W, Haluck-Kangas A, Gao QQ, Sarshad AA, Bartom ET, Stults A, Qadir AS, Hafner M and Peter ME (2018) CD95/Fas ligand mRNA is toxic to cells. eLife 7:e38621.

19. Rybak-Wolf A, Jens M, Murakawa Y, Herzog M, Landthaler M and Rajewsky N (2014) A variety of dicer substrates in human and C. elegans. Cell 159:1153–1167. doi: 10.1016/j.cell.2014.10.040

20. Luo QJ, Zhang J, Li P, Wang Q, Zhang Y, Roy-Chaudhuri B, Xu J, Kay MA and Zhang QC (2021) RNA structure probing reveals the structural basis of Dicer binding and cleavage. Nat Commun 12:3397. doi: 10.1038/s41467-021-23607-w

21. Cole C, Sobala A, Lu C, Thatcher SR, Bowman A, Brown JW, Green PJ, Barton GJ and Hutvagner G (2009) Filtering of deep sequencing data reveals the existence of abundant Dicer-dependent small RNAs derived from tRNAs. RNA 15:2147–60. doi: 10.1261/rna.1738409

22. Patel M, Bartom ET, Paudel B, Kocherginsky M, O’Shea KL, Murmann AE and Peter ME (2022) Identification of the toxic 6mer seed consensus for human cancer cells. Sci Rep 12:5130. doi: 10.1038/s41598-022-09051-w

23. Landthaler M, Gaidatzis D, Rothballer A, Chen PY, Soll SJ, Dinic L, Ojo T, Hafner M, Zavolan M and Tuschl T (2008) Molecular characterization of human Argonaute-containing ribonucleoprotein complexes and their bound target mRNAs. RNA 14:2580–96. doi: 10.1261/rna.1351608

24. Wang D, Zhang Z, O’Loughlin E, Lee T, Houel S, O’Carroll D, Tarakhovsky A, Ahn NG and Yi R (2012) Quantitative functions of Argonaute proteins in mammalian development. Genes Dev 26:693–704. doi: 10.1101/gad.182758.111

25. Hauptmann J, Kater L, Loffler P, Merkl R and Meister G (2014) Generation of catalytic human Ago4 identifies structural elements important for RNA cleavage. RNA 20:1532–8. doi: 10.1261/rna.045203.114

26. Liu J, Carmell MA, Rivas FV, Marsden CG, Thomson JM, Song JJ, Hammond SM, Joshua-Tor L and Hannon GJ (2004) Argonaute2 is the catalytic engine of mammalian RNAi. Science 305:1437–41.

27. Meister G, Landthaler M, Patkaniowska A, Dorsett Y, Teng G and Tuschl T (2004) Human Argonaute2 mediates RNA cleavage targeted by miRNAs and siRNAs. Mol Cell 15:185–97. doi: 10.1016/j.molcel.2004.07.007

28. Chu Y, Kilikevicius A, Liu J, Johnson KC, Yokota S and Corey DR (2020) Argonaute binding within 3’-untranslated regions poorly predicts gene repression. Nucleic Acids Res 48:7439–7453. doi: 10.1093/nar/gkaa478

29. Popp MW and Maquat LE (2018) Nonsense-mediated mRNA Decay and Cancer. Curr Opin Genet Dev 48:44–50. doi: 10.1016/j.gde.2017.10.007

30. Bartom ET, Kocherginsky M, Baudel B, Vaidyanathan A, Haluck-Kangas A, Patel M, O’Shea K, Murmann AE and Peter ME (2021) SPOROS: A pipeline to analyze DISE/6mer seed toxicity. PLOS Comp Biol 18:e1010022.

31. Su H, Trombly MI, Chen J and Wang X (2009) Essential and overlapping functions for mammalian Argonautes in microRNA silencing. Genes Dev 23:304–17. doi: 10.1101/gad.1749809

32. Houseley J and Tollervey D (2009) The many pathways of RNA degradation. Cell 136:763–76. doi: 10.1016/j.cell.2009.01.019

33. Ibrahim F, Maragkakis M, Alexiou P and Mourelatos Z (2018) Ribothrypsis, a novel process of canonical mRNA decay, mediates ribosome-phased mRNA endonucleolysis. Nat Struct Mol Biol 25:302–310. doi: 10.1038/s41594-018-0042-8

34. Glover ML, Burroughs AM, Monem PC, Egelhofer TA, Pule MN, Aravind L and Arribere JA (2020) NONU-1 Encodes a Conserved Endonuclease Required for mRNA Translation Surveillance. Cell Reports 30:4321–4331.e4. doi: https://doi.org/10.1016/j.celrep.2020.03.023

35. D’Orazio KN, Wu CC, Sinha N, Loll-Krippleber R, Brown GW and Green R (2019) The endonuclease Cue2 cleaves mRNAs at stalled ribosomes during No Go Decay. Elife 8. doi: 10.7554/eLife.49117

36. Tuck AC, Rankova A, Arpat AB, Liechti LA, Hess D, Iesmantavicius V, Castelo-Szekely V, Gatfield D and Bühler M (2020) Mammalian RNA Decay Pathways Are Highly Specialized and Widely Linked to Translation. Molecular Cell 77:1222–1236.e13. doi: 10.1016/j.molcel.2020.01.007

37. Behm-Ansmant I, Rehwinkel J, Doerks T, Stark A, Bork P and Izaurralde E (2006) mRNA degradation by miRNAs and GW182 requires both CCR4:NOT deadenylase and DCP1:DCP2 decapping complexes. Genes Dev 20:1885–98. doi: 10.1101/gad.1424106

38. Rehwinkel J, Behm-Ansmant I, Gatfield D and Izaurralde E (2005) A crucial role for GW182 and the DCP1:DCP2 decapping complex in miRNA-mediated gene silencing. RNA 11:1640–7. doi: 10.1261/rna.2191905

39. Hashimoto Y, Takahashi M, Sakota E and Nakamura Y (2017) Nonstop-mRNA decay machinery is involved in the clearance of mRNA 5’-fragments produced by RNAi and NMD in Drosophila melanogaster cells. Biochem Biophys Res Commun 484:1–7. doi: 10.1016/j.bbrc.2017.01.092

40. Orban TI and Izaurralde E (2005) Decay of mRNAs targeted by RISC requires XRN1, the Ski complex, and the exosome. RNA 11:459–69. doi: 10.1261/rna.7231505

41. Lima WF, De Hoyos CL, Liang XH and Crooke ST (2016) RNA cleavage products generated by antisense oligonucleotides and siRNAs are processed by the RNA surveillance machinery. Nucleic Acids Res 44:3351–63. doi: 10.1093/nar/gkw065

42. Pule MN, Glover ML, Fire AZ and Arribere JA (2019) Ribosome clearance during RNA interference. RNA 25:963–974. doi: 10.1261/rna.070813.119

43. Algeciras-Schimnich A, Pietras EM, Barnhart BC, Legembre P, Vijayan S, Holbeck SL and Peter ME (2003) Two CD95 tumor classes with different sensitivities to antitumor drugs. Proc Natl Acad Sci U S A 100:11445–50.

44. Golden RJ, Chen B, Li T, Braun J, Manjunath H, Chen X, Wu J, Schmid V, Chang TC, Kopp F, Ramirez-Martinez A, Tagliabracci VS, Chen ZJ, Xie Y and Mendell JT (2017) An Argonaute phosphorylation cycle promotes microRNA-mediated silencing. Nature 542:197–202. doi: 10.1038/nature21025

45. Hadji A, Ceppi P, Murmann AE, Brockway S, Pattanayak A, Bhinder B, Hau A, De Chant S, Parimi V, Kolesza P, Richards J, Chandel N, Djaballah H and Peter ME (2014) Death induced by CD95 or CD95 ligand elimination. Cell Rep 7:208–22. doi: 10.1016/j.celrep.2014.02.035

46. Hauptmann J, Schraivogel D, Bruckmann A, Manickavel S, Jakob L, Eichner N, Pfaff J, Urban M, Sprunck S, Hafner M, Tuschl T, Deutzmann R and Meister G (2015) Biochemical isolation of Argonaute protein complexes by Ago-APP. Proc Natl Acad Sci U S A 112:11841–5. doi: 10.1073/pnas.1506116112

47. Hafner M, Renwick N, Farazi TA, Mihailovic A, Pena JT and Tuschl T (2012) Barcoded cDNA library preparation for small RNA profiling by next-generation sequencing. Methods 58:164–70. doi: 10.1016/j.ymeth.2012.07.030

48. Lorenz R, Bernhart SH, Honer Zu Siederdissen C, Tafer H, Flamm C, Stadler PF and Hofacker IL (2011) ViennaRNA Package 2.0. Algorithms Mol Biol 6:26. doi: 10.1186/1748-7188-6-26

49. Hulsen T, de Vlieg J and Alkema W (2008) BioVenn - a web application for the comparison and visualization of biological lists using area-proportional Venn diagrams. BMC Genomics 9:488. doi: 10.1186/1471-2164-9-488

50. Huang da W, Sherman BT and Lempicki RA (2009) Systematic and integrative analysis of large gene lists using DAVID bioinformatics resources. Nat Protoc 4:44–57. doi: 10.1038/nprot.2008.211

51. Sherman BT, Hao M, Qiu J, Jiao X, Baseler MW, Lane HC, Imamichi T and Chang W (2022) DAVID: a web server for functional enrichment analysis and functional annotation of gene lists (2021 update). Nucleic Acids Res. doi: 10.1093/nar/gkac194

