## Supplementary figures for "CD95/Fas ligand mRNA is toxic to cells through more than one mechanism"

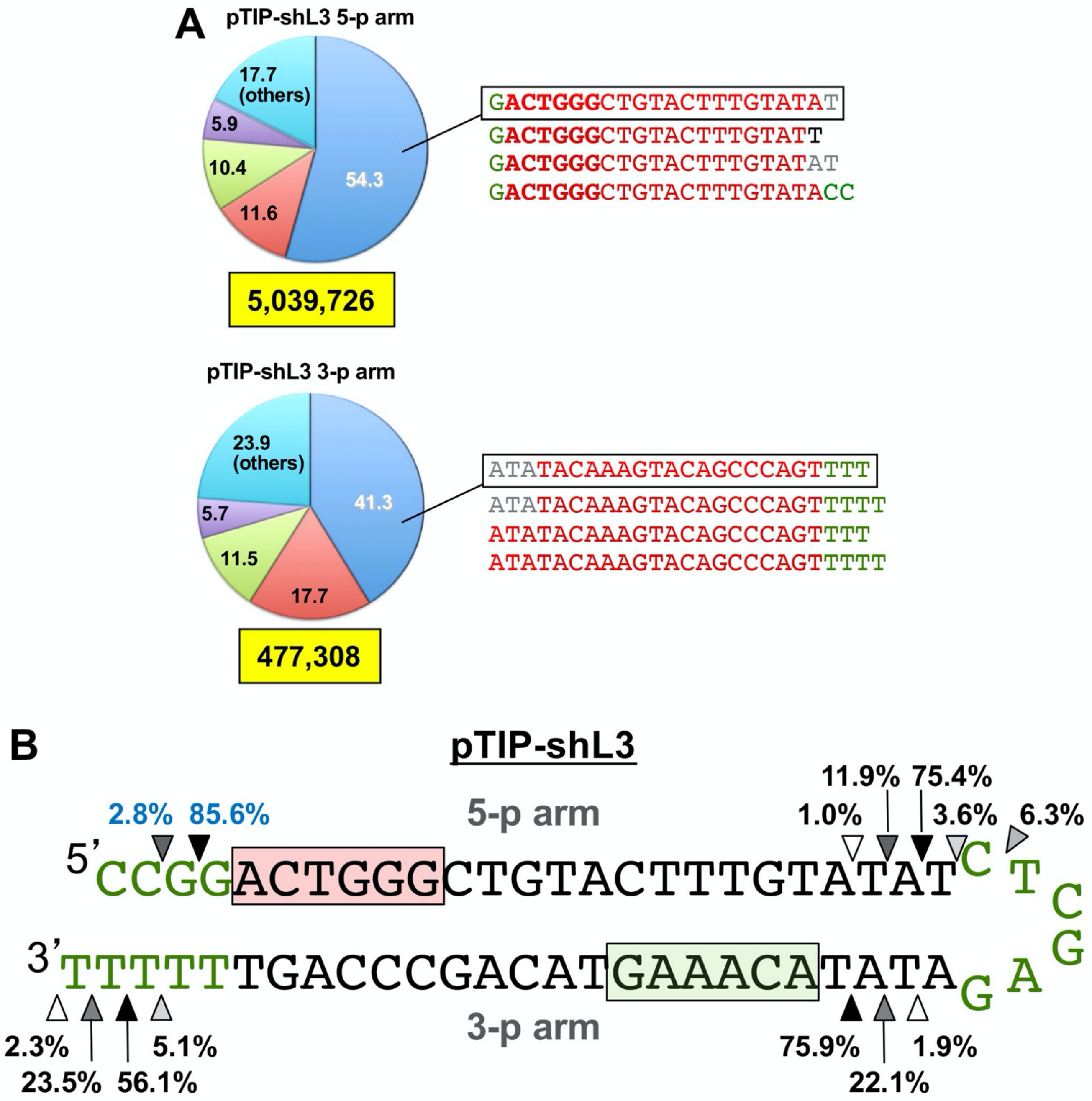


**Figure S1 - Processing of pTIP-shL3 results in preferential selection of the sense strand of the shRNA**
Reads mapping to shL3 in 293T stably expressing pTIP-shL3 50 hours after dox induction. **(A)** Pie charts representing the contribution of each shL3-derived sequence to the total reads derived from the two shRNA arms. Sequences derived from the pTIP-shL3 5-parm (sense strand, top) and the 3-p arm (antisense strand, bottom) are shown. The top four most abundant sequences derived from each arm are listed. Nucleotides shown in green are derived from the viral vector, in red are shRNA sequences, in grey are missing nucleotides, and in black are additional nucleotides. **(B)** Schematic representing the secondary structure of shL3. The frequency of each cleavage event in the RNA-Seq data and the site of cleavage is indicated with arrows. The most common cleavage site is indicated with a black arrow, and the least common with a white arrow. Nucleotides derived from the pTIP shRNA vector are in green. The 6mer seed of the most abundant generated shRNA on the 5-p arm is indicated in the red box, the one on the 3-p arm in a green box.

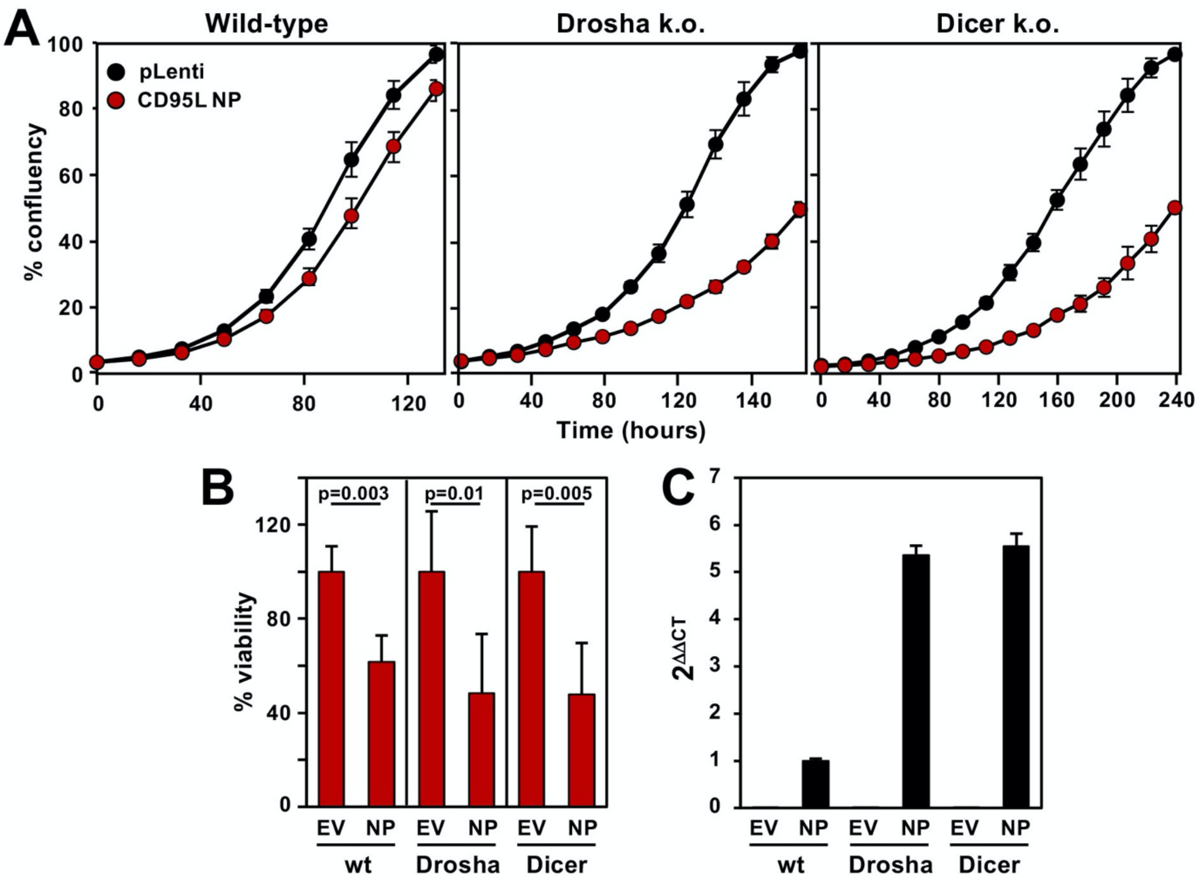


**Figure S2 - Dicer is not required for pLenti CD95L NP toxicity**
**(A)** Percent confluency over time of HCT116 wt (left) HCT116 Drosha k.o. (center) and HCT116 Dicer k.o. cells (right) expressing either pLenti or pLenti-CD95L NP. Data are representative of >3 independent experiments. **(B)** Percent viability of the cells in A 120 hrs after infection. Viability of CD95L expressing cells was normalized to pLenti (EV) expressing cells of the same genotype. Error bars represent the standard deviation (SD) of three replicates. (T-test, Benjamini corrected p-value). **(C)** Real-time qPCR assessment of CD95L mRNA expression in HCT116 cells of each genotype. Expression was normalized to HCT116 cells expressing pLenti-CD95L NP. Error bars represent the SD of sample triplicates. Data representative of >3 independent experiments.

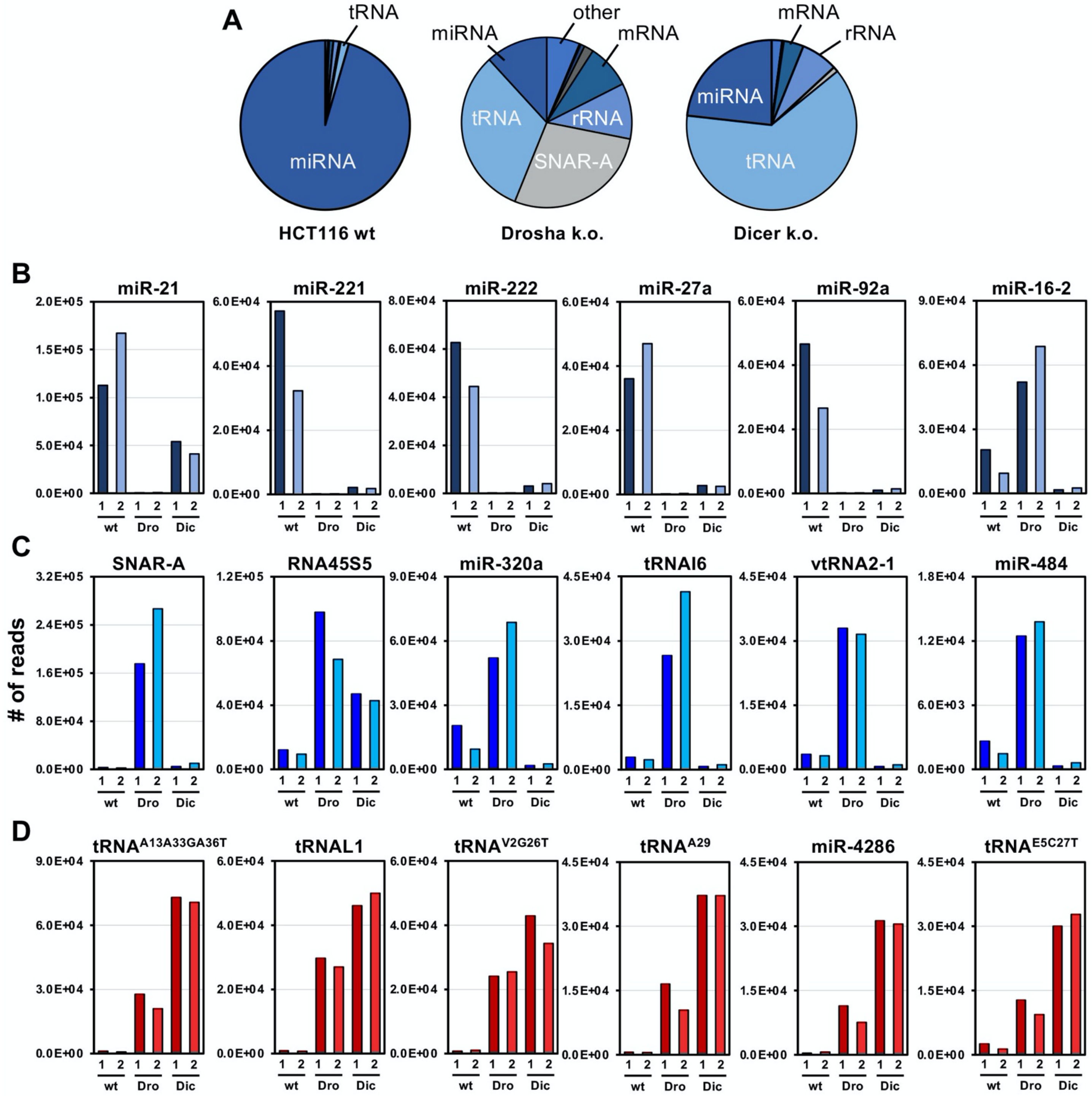


**Figure S3 - Characterization of R-sRNAs in cells lacking Dicer and Drosha**
**(A)** Pie charts representing the types of sRNAs that occupy the RISC of HCT116 cells by genotype. **(B)** Number of reads normalized per million of the top five most abundant sRNAs in the RISC of HCT116 cells (navy), **(C)** HCT116 Drosha k.o. cells (blue), and **(D)** HCT116 Dicer k.o. cells (red). Each bar represents read counts in one sample replicate, sample replicate 1 (darker shade) and replicate 2 (lighter shade).

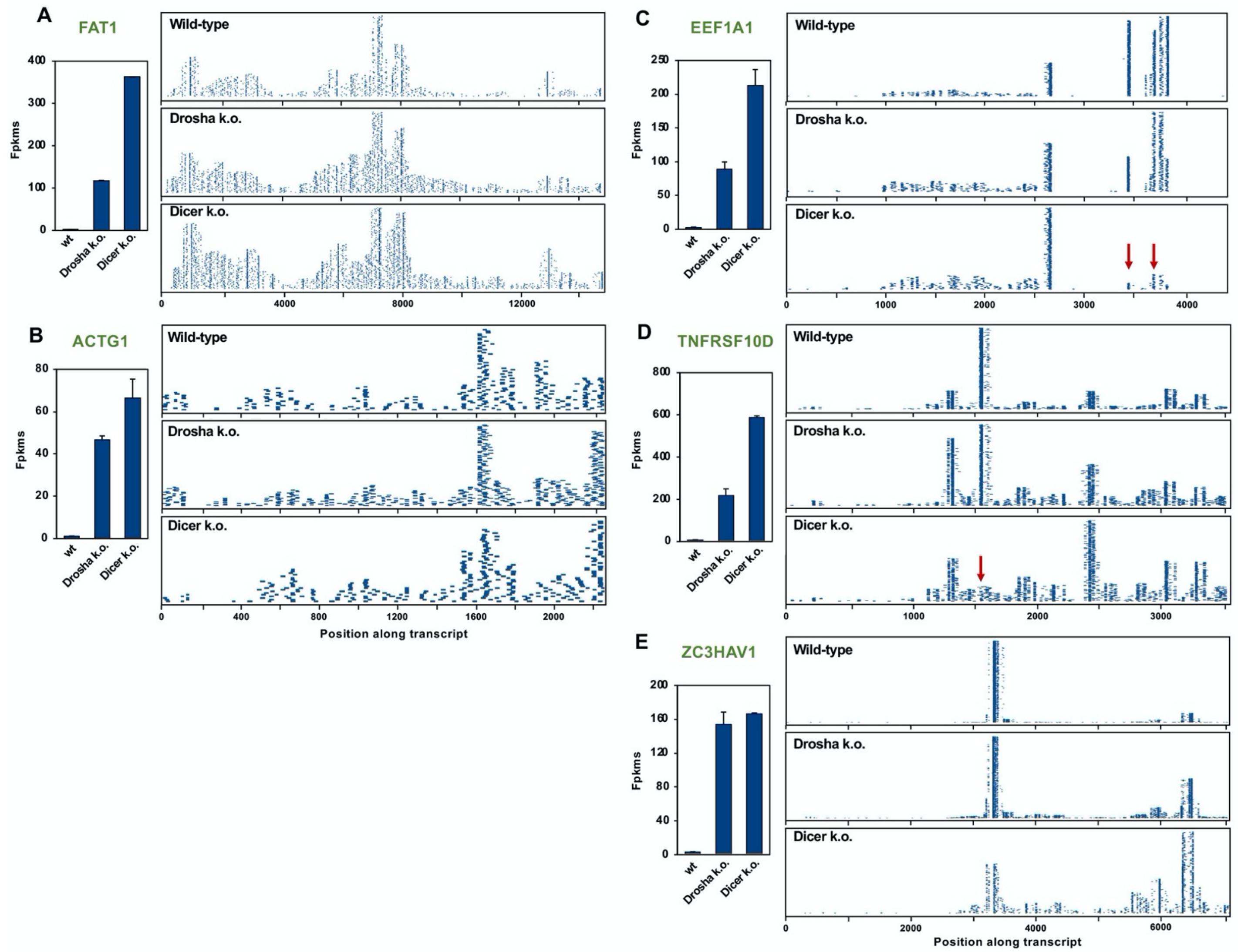

**Figure S4 - Endogenous mRNAs are processed and loaded into the RISC of Dicer k.o. cells**
**(A-E)** Mapping of R-sRNAs to the transcripts of five selected highly processed mRNAs in cells infected with pLenti-CD95L NP. One horizontal line represents one read in the RISC of HCT116 (top), Drosha k.o. (center) and Dicer k.o. cells (bottom). *Left*, Reads from both replicates are combined and error bars show variance. *Right*, normalized read counts (fpkms) by genotype. Selected processed mRNAs **(A)** FAT1, **(B),** ACTG1, **(C)** EEF1A1, **(D)** TNFRSF10D, and **(E)** ZC3HAV1. Red arrows indicate stacks with few reads in Dicer k.o. cells.

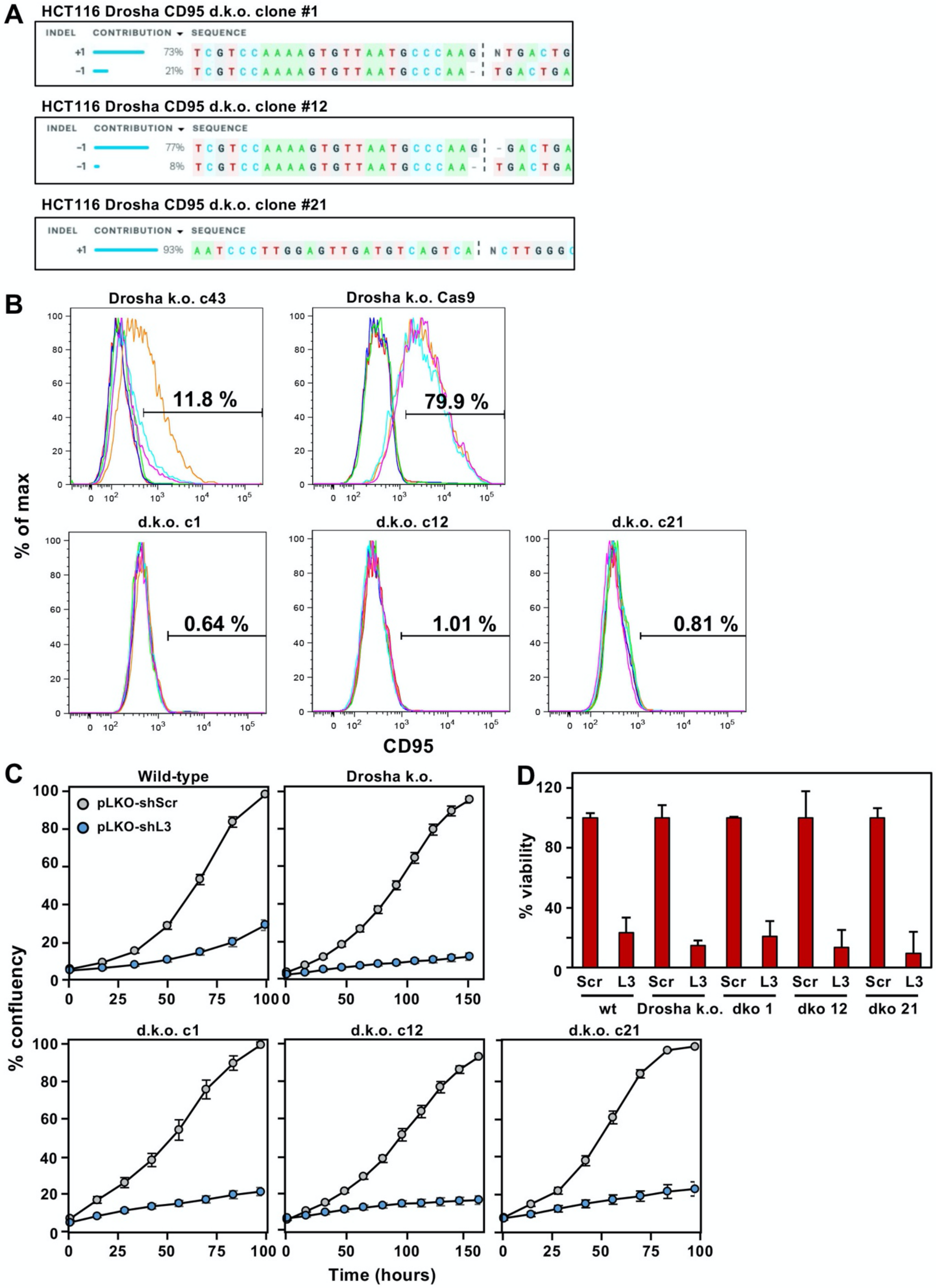


**Figure S5 - Characterization of HCT116 Drosha CD95 d.k.o. cells**

(**A**) Representation and contribution of indel mutations in the DNA of single cell clones analyzed by Synthego ICE tool. **(B)** Surface staining of HCT116 and Drosha CD95 d.k.o. clones for CD95. Gates represent % positivity. (**C**) Cell confluency over time in HCT116 cells expressing either pLKO-shL3 or pLKO-shScr. HCT116 , Drosha k.o., and d.k.o. cells were assayed. Bars indicate standard error of triplicates. **(D)** Relative cell viability at 96 hours. Error bars represent standard deviation of triplicates.

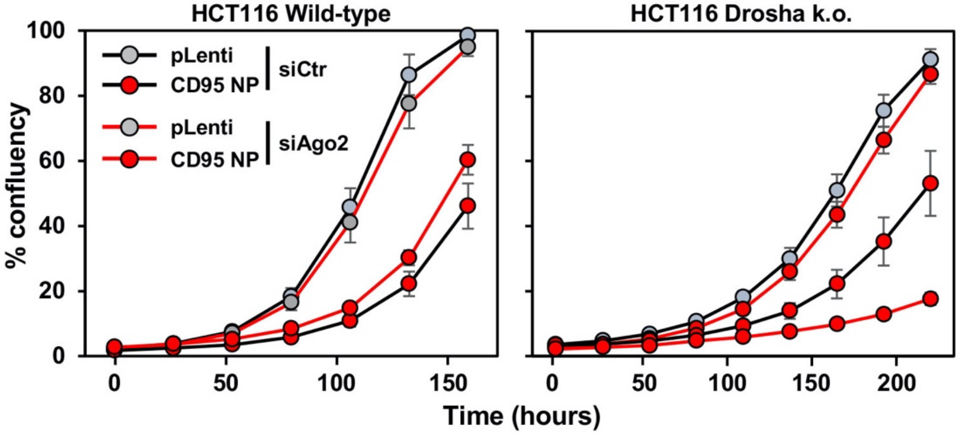


**Figure S6** - **The role of Ago2 in mediating CD95L toxicity is cell type specific**
Percent cell confluency over time in HCT116 (left) and HCT116 Drosha k.o. (right) cells transfected with 25 nM siAgo2 or siCtr and subsequently infected with pLenti or pLenti-CD95L NP. Data is representative of two independent experiments.

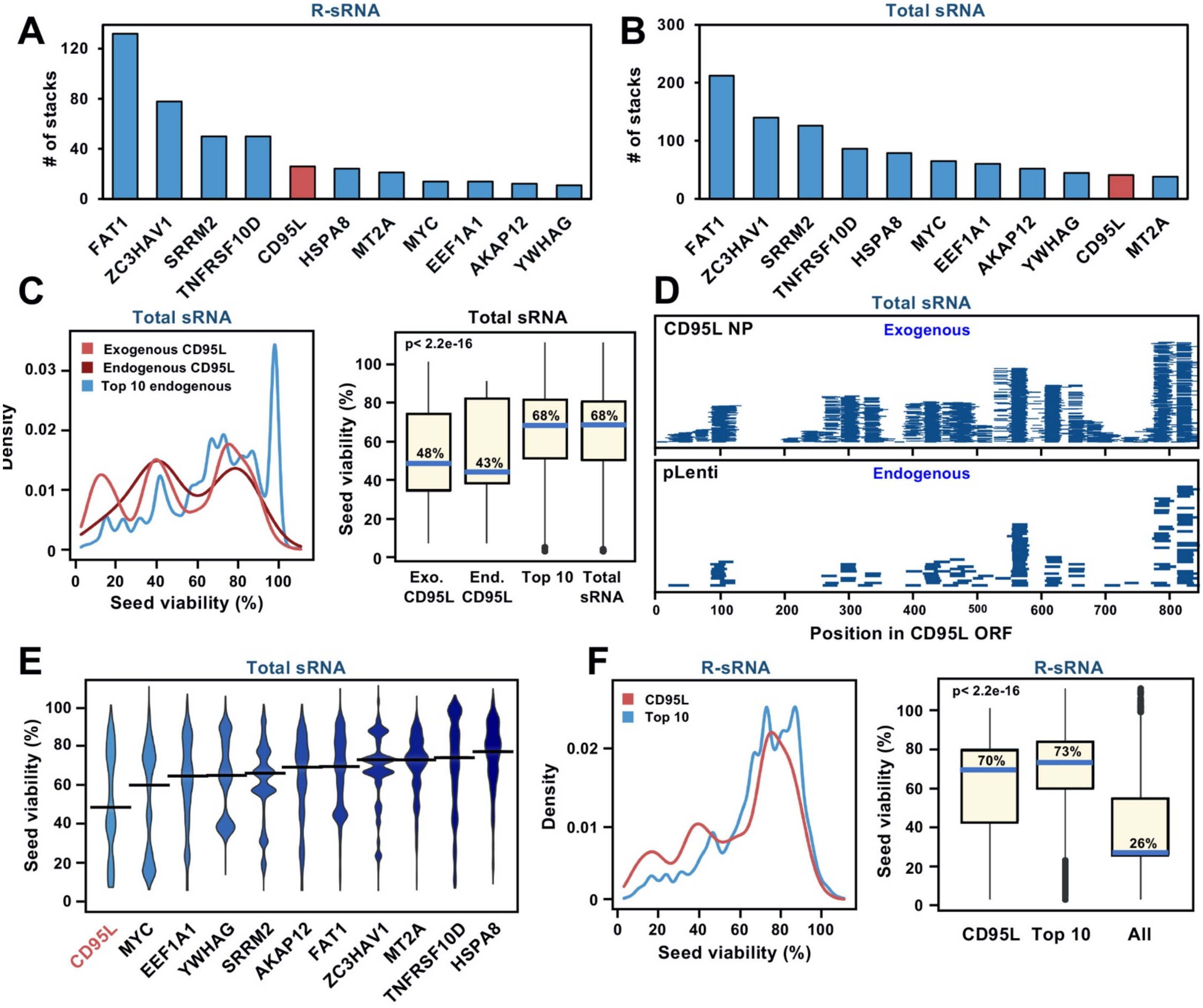


**Figure S7 - CD95L-derived reads skew more toxic than reads derived from other mRNAs**

**(A, B)** Ranking of highly expressed and processed protein coding genes by number of stacks. A stack was defined as reads, 10 or more, with the same 5' start site. pLenti-CD95L NP is indicated in red. Ago bound reads are represented in (**A)** and stacks in the total sRNA are represented in (B)**.** Reads from two replicates were co0mbined. **(C)** *Left*, Density plot representing the predicted 6mer seed viability of reads 18-25 nt long derived from CD95L vs the Top 10 most abundant and highly processed mRNAs in the total sRNA in aggregate. *Right*, box plots representing the skew of the 6mer seed viability of the derived reads. Blue lines indicate the median 6mer seed viability. Kruskal-Wallis p-value < 2 x $10^{-16}$. **(D)** Mapping of CD95L-derived sRNAs found in the total sRNA to the CD95L ORF. Reads derived from exogenously expressed CD95L (top), and endogenous CD95L-derived reads (bottom). **(E)** Violin plots representing the distribution and the median (black solid line) 6mer seed viability of each of the Top 10 processed mRNAs and exogenously expressed CD95L (red). **(F)** *Left*, Density plot representing the predicted 6mer seed viability of R-sRNAs derived from CD95L vs the Top 10 processed mRNAs. Endogenous CD95L reads were too few to plot. *Right*, box plots representing the skew of the 6mer seed viability of R-sRNAs derived from CD95L, the Top 10 processed mRNAs and the total RISC content (Kruskal-Wallis p-value < 2 x$10^{-16}$).

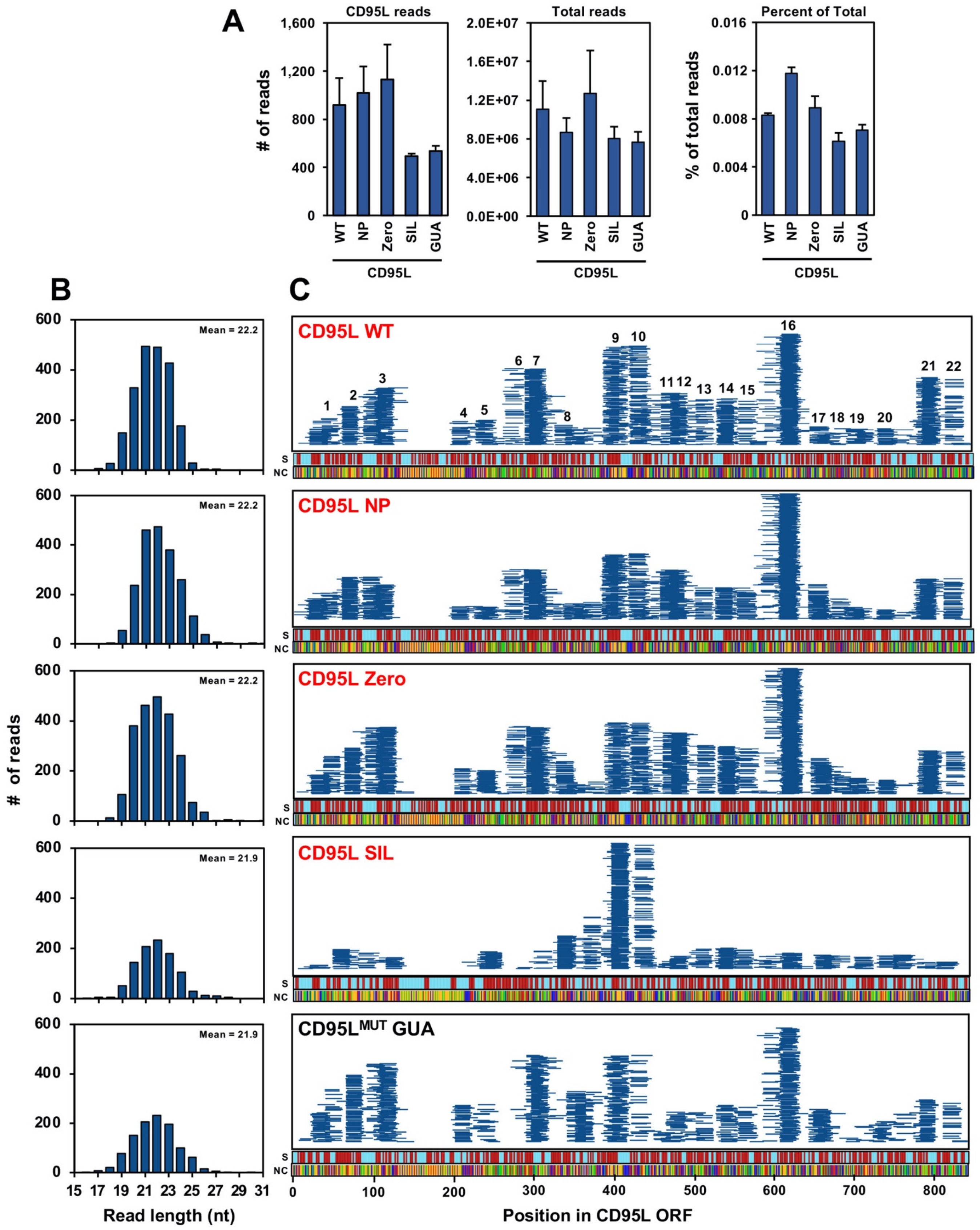


**Figure S8 - Processing and RISC loading of CD95L mutant derived reads**

RISC bound reads of CD95L mutants expressed in CD95 d.k.o. 12 cells were analyzed. **(A)** *Left*, raw counts of reads aligning to CD95L mutant sequences. *Center*, total raw reads sequenced per sample. *Right*, percentage of total raw reads that are derived from pLenti-CD95L mutants. The average of two replicates is shown. Error bars represent the variance of the mean. **(B)** Bar plots representing the read lengths of various CD95L derived reads in the RISC (Kruskal-Wallis p-value was determined). **(C)** Mapping of CD95L-derived sRNAs along the ORF of each CD95L mutant. Each horizontal line represents one read. Reads from both replicates are displayed. Toxic mutants are labeled in red and non-toxic mutants in black. Beneath each stack plot, stem-loop regions (S) are mapped with stems (red) and loops (blue). The next bar below represents the nucleotide content of each CD95L mutant sequence, **A**denine (blue), **U**racil (green), **G**uanine (red), and **C**ytosine (yellow).
